# New molecular signatures defining the differential proteostasis response in ALS-resistant and -sensitive motor neurons

**DOI:** 10.1101/2022.04.10.487765

**Authors:** Ana Paula Zen Petisco Fiore, Shuvadeep Maity, Disi An, Justin Rendleman, Dylan Iannitelli, Hyungwon Choi, Esteban Mazzoni, Christine Vogel

## Abstract

Amyotrophic Lateral Sclerosis (ALS) is a fatal adult neurodegenerative disease characterized by proteostasis dysregulation, resulting in progressive loss of spinal and upper motor neurons. A subset of cranial motor neurons resistant to ALS-stress survive until late stages of the disease. To investigate these differences, we exploited a unique platform of induced cranial and spinal motor neurons (iCrMNs and iSpMNs, respectively). Exposing both cell types to proteotoxic stress, we quantified transcriptome and proteome changes over 36 hours for a core set of >8,200 genes. While mRNA and protein changes under stress were congruent for many genes, cell-type specific differences manifested at either the RNA or protein level, but less at both. At the protein level, iCrMNs and iSpMNs differed significantly with respect to abundance of many membrane proteins, including synaptic proteins, solute carriers, adhesion molecules, and signaling molecules suggesting that the superior stress survival of iCrMNs involve diverse pathways supporting neuronal function. Other differences included genes involved in ribosome biogenesis and subunits of the core proteasome. We investigated the role of proteasomal degradation in more detail. Our data showed that although stress reduces proteasome activity in both neuronal types, iCrMNs had significantly more abundant and active 26S proteasome than iSpMNs, which indicate a higher capacity for the degradation of ubiquitinated proteins. We identified a new regulator of this better performance, i.e. the nuclear proteasome activator Ublcp1, whose inhibition sensitized iCrMNs, but not iSpMNs, to stress and abolished their higher survival rates. The results suggest that the two neuronal cell types regulate and use the degradation machinery differently under normal and stress conditions. Overall, this work demonstrates the value of unbiased system-wide analyses in generating hypotheses on differential proteostasis regulation in cranial and spinal motor neurons.

## Introduction

Neurodegenerative diseases are characterized by progressive deterioration of specific neuron populations, while other neuronal cell types are resistant ^1–3^. For example, during Amyotrophic Lateral Sclerosis (ALS), most spinal motor neurons progressively degenerate leading to muscle denervation and eventual paralysis and death by respiratory failure ^4^. In contrast, a subset of cranial motor neurons (oculomotor, trochlear, and abducens motor neurons) typically survive and remain functional until late stages of the disease ^5–7^. While mutations in several genes - such as SOD1, TDP43 or C9ORF72 - have been linked to the manifestation of familial ALS, the molecular mechanisms behind sporadic ALS, which involves >90% of the patients, are not understood: it is not known if and which additional proteins and pathways might substantiate the differences between cranial and spinal motor neurons. The reasons for the differential ALS-sensitivity of the two motor neuron types are unknown, and systematic comparison of the two cell types has been largely hindered by the lack of suitable experimental systems.

We exploited a unique cell model that we recently developed to show that induced spinal and cranial motor neurons (iSpMNs, iCrMNs) derived from mouse embryonic stem cells differ significantly in their resistance to protein misfolding stress ^8^. The accumulation of misfolded proteins resulted from altered proteostasis regulation and is a common feature of ALS ^9–12,13–16^. We showed that part of the iCrMNs’ superior stress resistance was due to their heightened ability to ubiquitinate and degrade proteins ^8^.

However, the response to proteotoxic stress involves a variety of pathways, ranging from the activation of survival pathways to the initiation of cell death ^17–19^ - and the differential role of these pathways in the induced cranial and spinal motor neurons is not known. Further, chemical and genetic activation of the proteasome in iSpMNs could only partially rescue their survival in response to proteotoxic stress ^8^, suggesting the role of additional pathways in the cells’ differential proteostasis control. These findings are consistent with what is known about ALS in general: individual genes and pathways can only explain a small fraction of disease cases, and we have no understanding of the molecular signatures that define ALS-sensitive and -resistant cell types.

To identify differences that could explain the differential neuronal response to proteotoxic stress, we devised a study in which we created an unbiased, large-scale transcriptome and proteome dataset of >8,200 genes, monitoring iSpMNs and iCrMNs responding to protein misfolding stress over 36 hours. We used our expertise in integrative data analysis ^20^ to extract significant differences between the motor neuron types, both under normal and stress conditions. We investigated one of these differences in more detail, elucidating aspects of differential regulation of the nuclear proteasome in the motor neurons.

## Results

### Dynamic profiling of the response to misfolded protein stress of motor neurons

We generated highly pure populations of induced cranial and spinal motor neurons from an inducible mouse stem cell system described previously ^8,21^. Mirroring our prior observations ^8^ the cells showed significant sensitivity to protein misfolding stress induced by cyclopiazonic acid (CPA). We devised an experimental setup to collect transcriptomic and proteomic data for >13,000 and >8,600 transcripts and proteins, respectively (**Figure 1A, Supplementary Datasets 1** and **2**). We examined 3 time points (0, 12, and 36 hours) in two independent replicates and included a control experiment with cells treated with Dimethyl sulfoxide (DMSO). We chose the CPA concentration and time points to focus on the stress response and avoid extensive apoptosis. The conditions correctly induced phosphorylation of PERK, a hallmark of the unfolded protein response (UPR)(**Supplementary Figure S1**). The measurements were highly reproducible between replicates (**Supplementary Figure S1**).

**Figure 1.**
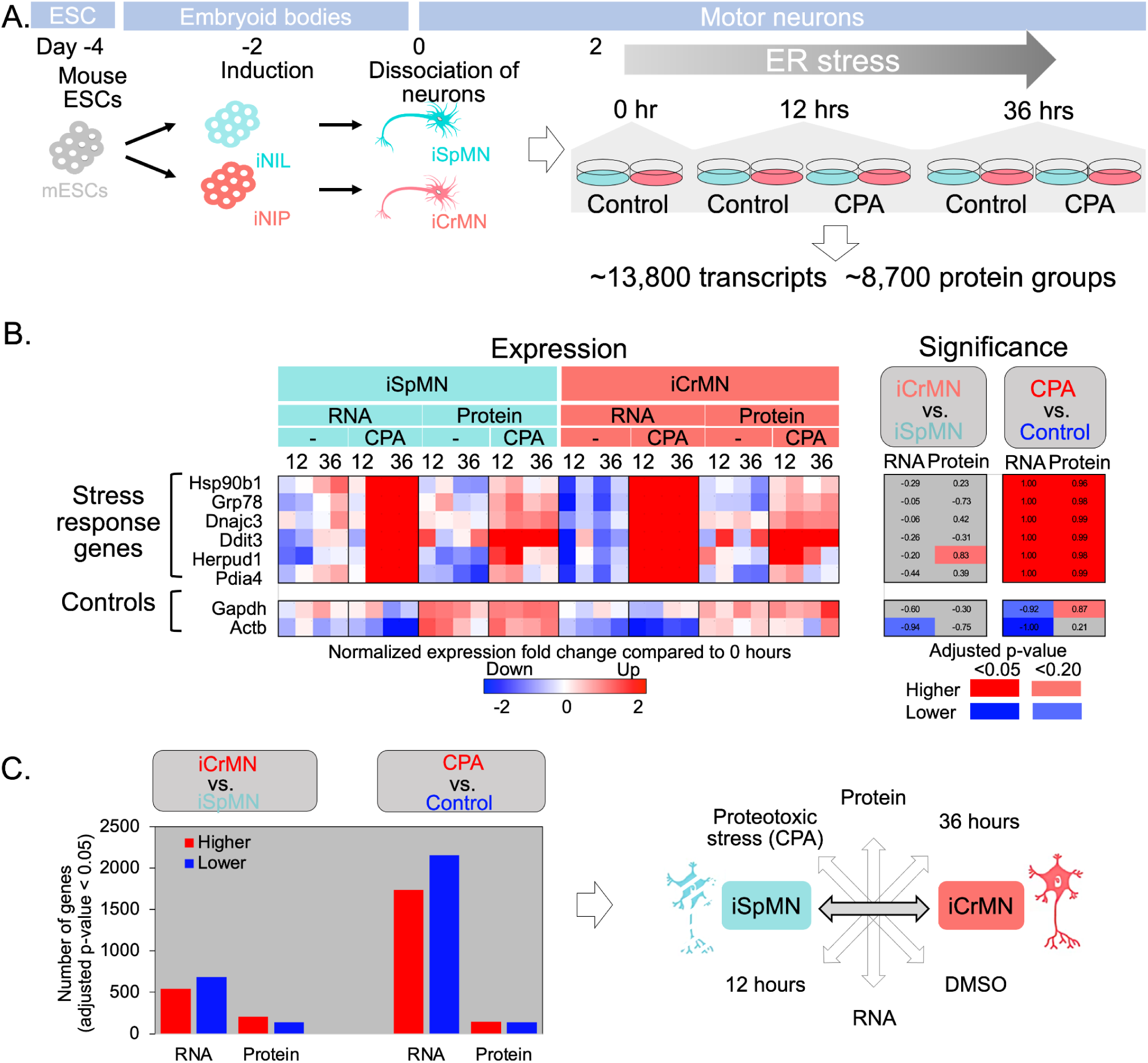
Multi-dimensional transcriptomics and proteomics of induced spinal and cranial motor neurons. **A.** We induced mouse ESC-derived cranial (iCrMN) and spinal (iSpMN) using a well established protocol ^8,21^. Two days after differentiation, cells were treated with cyclopiazonic acid (CPA) to induce protein misfolding and with DMSO as control. Samples were collected at 0, 12, and 36 hours post-treatment. The experiment was conducted in duplicates. We quantified ∼13,800 transcripts and ∼8,700 protein expression data, amounting to 8,206 genes with complete data. **B.** The panels show examples of stress response genes whose expression increases at both the RNA and protein levels. The heatmap on the left illustrates the expression data; values are Z-score normalized log-transformed ratios of the expression value at 12 or 36 hours compared to 0 hours. The heatmap on the right illustrates the outcomes of the significance analysis (ANOVA, adjusted p-values). Adjusted p-values are directed: shades of red if the direction of the expression difference is positive, shades of blue if the direction is negative. Adjusted p-values are also thresholded at 0.05 and 0.20 as shown in the legend. Stress-response genes are significantly upregulated at both the RNA and protein level (adjusted p-value < 0.05) while not different between the cell types. Control genes show no significant expression differences at the protein level for either cell type or stress (adjusted p-value < 0.05). **C.** Total numbers of genes with significant expression differences with respect to either RNA or protein expression levels testing the two cell types (left) and stress vs. control conditions (right)(adjusted p-value < 0.05). Red denotes a positive expression difference, blue denotes a negative expression difference. The largest number of significant regulatory events were observed for RNAs down-regulated in response to stress (n=2,154). CPA - cyclopiazonic acid; iSpMN - induced spinal motor neurons; iCrMN - induced cranial motor neurons

We proceeded to construct a core dataset comprising 8,206 genes with complete RNA and protein information and conducted several tests to confirm the dataset’s quality (**Supplementary Figure S1**). **Supplementary Dataset S1** shows the extended dataset at the RNA level only. We analyzed the core dataset with respect to differences between cell types, conditions, and time points using 3-way ANOVA and heatmaps to evaluate and visualize the statistical significance and abundance values, respectively. Across all figures, heatmaps display *changes* in expression levels, i.e. values were normalized to those from the first time point. In comparison, western blots and activities show levels in comparison to the loading control. ANOVA was conducted on the individual (Cell-type, Stress, Time) and on the interaction denominators (Cell type/Stress, Cell-type/Time, Stress/Time, Cell type/Stress/Time). Only Cell-type and Stress and Time resulted in significant changes (adjusted p-value < 0.05).

To illustrate the outcomes of these analyses, we show known UPR genes and controls (Gapdh, Actb) in **Figure 1B**, with the expression patterns on the left and the significance values on the right. We visualized the results of the significance analysis using dark and light colors for differences at an adjusted p-value cutoff of <0.05 and <0.20, respectively. Positive differences, e.g. higher abundance in iCrMN compared to iSpMN or higher abundance under stress compared to control, were marked in shades of red; negative differences in shades of blue. Correspondingly, all stress marker genes showed significant induction at both the RNA and protein level (adjusted p-value < 0.05), but no substantial differences between the two motor neuron types, further confirming that the experiment correctly induced proteotoxic stress in both cell types.

**Figure 1C** provides an overview of the statistically significant differences identified in the core dataset (adjusted p-value<0.05; **Supplementary Dataset S2**). Most of the significant changes occurred in response to protein misfolding stress, in particular at the RNA level. **Supplementary Figure S2** shows the expanded results for all denominators. The interaction terms, e.g. Cell type/Stress, did not result in any significant differences at the 0.05 cutoff. We also observed several hundreds of significant differences between the two motor neuron types, at both the RNA and protein level, regardless of the genes’ response to stress (adjusted p-value < 0.05). The remaining analysis focused on this set of genes to further our understanding of the molecular characteristics specific to iCrMNs and iSpMNs with the goal of testing new hypotheses on their differential ALS sensitivity.

### Complex molecular signatures marking differences between cell types

**Figure 2** provides an overview of the core dataset, illustrating the similarities between the two cell types and the complex expression dynamics in response to stress. The figure highlights differences between RNA and protein expression, suggesting an extensive regulation at both the transcriptional and post-transcriptional level. We grouped expression patterns into 25 clusters (**Figure 2A**) and visualized the outcomes of the significance analysis (ANOVA) for both cell type and stress (**Figure 2B**). We also analyzed the 25 clusters for enriched functions: indeed, we find several clusters with increased expression levels under stress (e.g. 4, 15, 16, and 17), and genes in these clusters are enriched for stress response pathways, further confirming the correct experimental design (**Supplementary Figure S3**).

**Figure 2.**
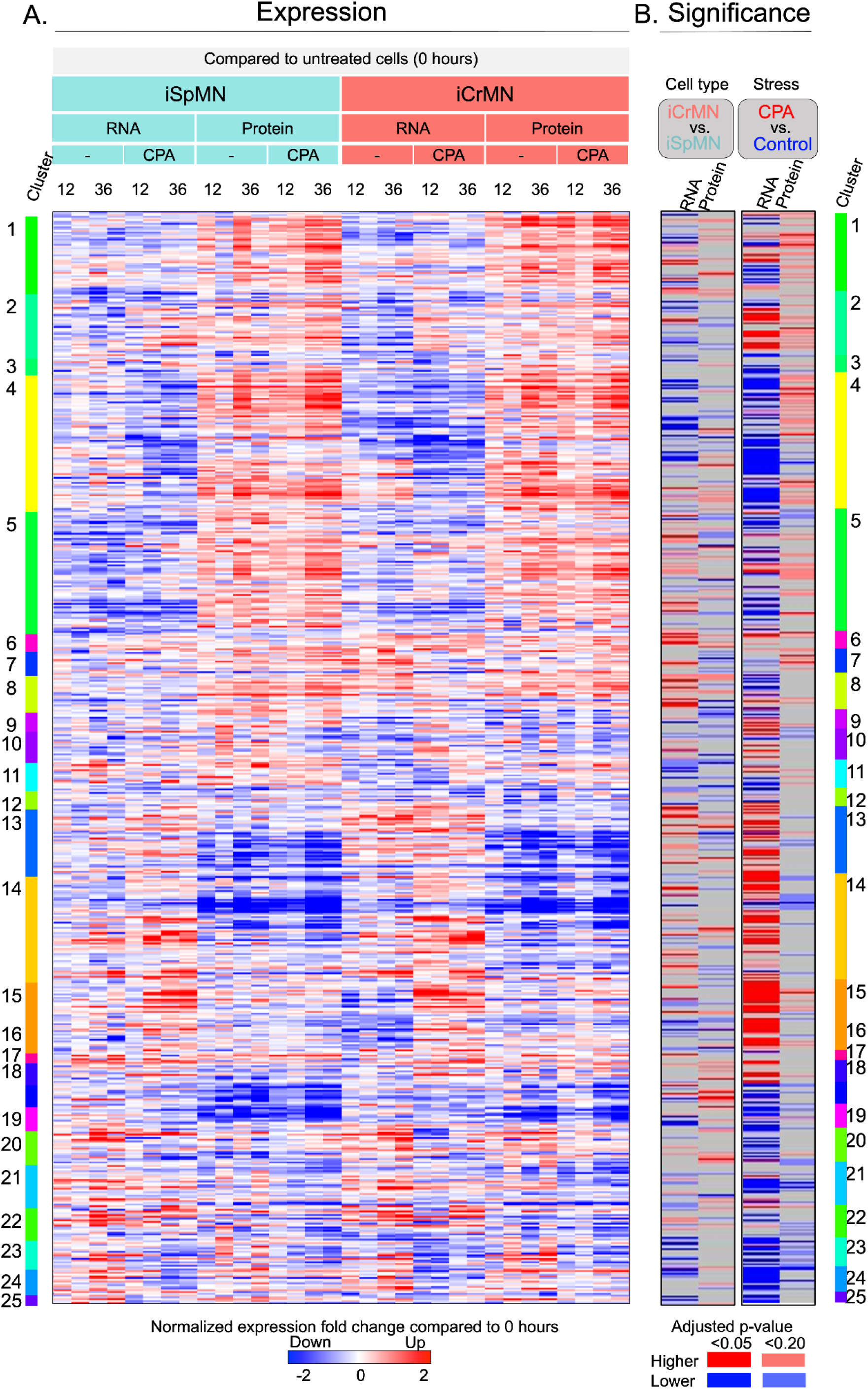
Complex transcriptome and proteome expression changes of spinal and cranial motor neurons responding to proteotoxic stress. **A.** Heatmap showing expression differences as Z-score normalized log-transformed ratios of the expression value at 12 or 36 hours compared to 0 hours of treatment with cyclopiazonic acid or DMSO, for the 8,206 genes with complete data. **B.** Heatmaps showing the significance values (adjusted p-values) from the ANOVA analysis for the differences between cell types and between stress and control conditions. Adjusted p-values were directed for better visualization of the results: shades of red if the direction of the expression difference is positive (enriched in iCrMN), shades of blue if the direction is negative (enriched in iSpMN). Adjusted p-values were also thresholded at 0.05 and 0.20 as shown in the legend. Grey indicates no significant difference. CPA - cyclopiazonic acid; iSpMN - induced spinal motor neurons; iCrMN - induced cranial motor neurons

Most of the significant cell type differences were specific to either the RNA or the protein level; only a small fraction of the observed differences were shared between protein and RNA (adjusted p-value<0.05, **Figure 3**, left). This result indicates that cell specificity might be implemented through either transcriptional and post-transcriptional regulation, but perhaps not both. The interpretation of this observation, i.e. why the cells would adjust mRNA levels but apparently not protein levels, is subject to future investigation. One possible technical explanation lies in the fact that protein changes are typically smaller in magnitude and therefore more difficult to detect with statistical confidence.

**Figure 3.**
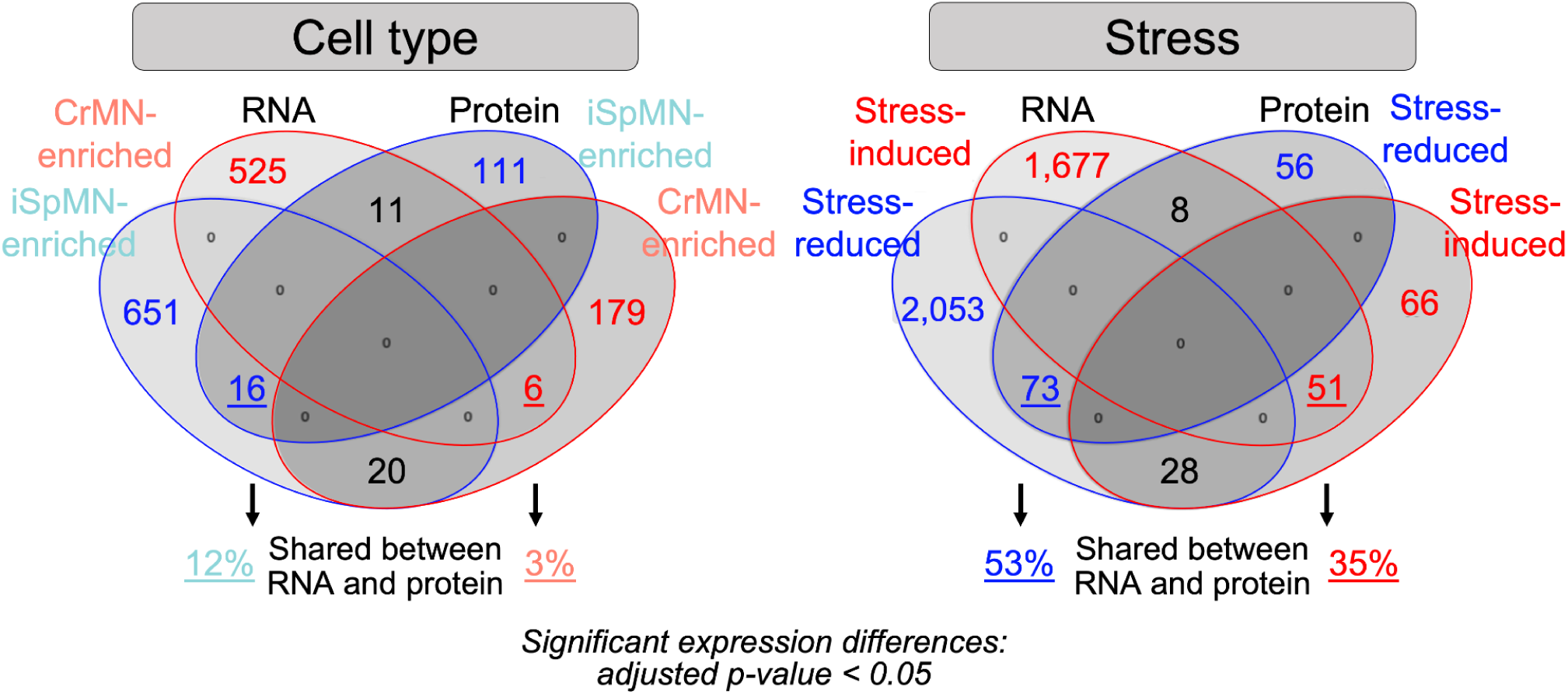
Shared and specific differences between cell types and in response to stress. The Venn diagrams summarize the numbers of differentially expressed transcripts and proteins between cell types (left) and stress and normal conditions (right)(ANOVA, adjusted p-value < 0.05), split into subsets according to the direction of the change. Most genes change specifically at either the RNA or protein level, but not both. CPA - cyclopiazonic acid; iSpMN - induced spinal motor neurons; iCrMN - induced cranial motor neurons

In comparison, a large proportion of the significant changes in response to misfolding stress occurred at both the RNA and protein level (53% and 35%, respectively; adjusted p-value<0.05, **Figure 3**, right), indicating that the response to stress in both cell types is highly coordinated across different levels of gene expression regulation. Fewer genes demonstrated RNA and protein changing in opposite directions: for example, one fifth of the significantly stress-induced proteins were significantly repressed at the transcript level (19%, 28/145). Such induction during stress is typical for the UPR: despite the global translation shutdown which is part of stress response, many stress response genes are translationally induced ^22,23^.

**Table 1** summarizes enriched functions amongst genes with significant differences at the RNA or protein level (FDR<0.05), as well as the fold-enrichment of the function compared to the background. Bold font indicates significant and major function categories; normal font indicates example pathways and genes (**Table 1**). iCrMNs were significantly enriched for membrane proteins and their corresponding transcripts including solute carriers, ion channels, synaptic proteins and G-proteins, proteins from the proteasome core particle and transcripts from Rap1 signaling (FDR<0.05, **Table 1**). iSpMNs were significantly enriched for proteins involved in ribosome biogenesis, and transcripts involved in the citric acid cycle, protein folding, membrane proteins, as well as of the regulatory particle of the proteasome (FDR<0.05, **Table 1**).

**Table 1.** Function enrichment amongst differentially expressed mRNAs and proteins. Summary of the significantly enriched functions and pathways amongst genes with significantly differential expression as shown in **Figure 1C** (ANOVA, adjusted p-value < 0.05). The function enrichment was conducted separately for RNAs and proteins with significantly higher (red) or lower (blue) expression in iCrMN/iSpMN or under stress/no stress, i.e. for eight sets of genes. Bold text indicates function categories enriched with a false discovery rate < 0.05; normal text indicates examples of genes and pathways within that category. Black bars with numbers indicate fold-enrichment of the function category (bold) compared to the background (8,206 genes). Gray background indicates function categories that are similar between the RNA and protein data.

| RNA |  | Protein |  |
| --- | --- | --- | --- |
| iCrMN enriched |  |  |  |
| Membrane proteins, e.g. ion transport, channels, cell adhesion | 1.4 | Membrane proteins, e.g solute carriers, synaptic proteins, G-proteins, receptors | 1.6 |
| Rap protein signal transduction, i.e. Rap1b, Rap2b, Rapgef1, Rapgef2, Plk2 | 9.3 | Proteasome core complex | 23.0 |
| iSpMN enriched |  |  |  |
| Proteasome regulatory particle | 8.3 | Ribosome biogenesis, RNA processing | 6.4 |
| Membrane proteins, e.g. GPI-anchored proteins, receptors | 6.4 |  |  |
| Citric acid cycle, Oxidative phosphorylation | 5.0 |  |  |
| Protein folding, e.g. chaperones | 2.2 |  |  |
| RNA |  | Protein |  |
| Stress-induced in both cell types |  |  |  |
| Response to endoplasmic reticulum stress | 3.2 | Response to endoplasmic reticulum stress | 7.8 |
| SRP-dependent cotranslational protein targeting to membrane, i.e. Sec63, Sec61a1, Srp9, Srp19, Srp68, Srp72, Srpr, Ssr3 | 4.2 | Endoplasmic reticulum | 2.5 |
| tRNA aminoacylation | 2.6 |  |  |
| Stress-reduced in both cell types |  |  |  |
| Ion transport | 1.4 | Ion transport., e.g. ion channels | 7.5 |
| Synapse, e.g. synaptic vesicle, nervous systems development | 1.8 | Nervous system development, e.g. cell differentiation | 4.4 |
| Cell differentiation | 1.4 | Growth cone | 4.1 |
| G-protein coupled receptor signaling | 1.7 | Membrane proteins, e.g. cell adhesion, ion channel | 1.8 |
| Cell adhesion | 2.5 |  |  |
| Cholesterol biosynthesis | 2.8 |  |  |
| Citric acid cycle, Oxidative phosphorylation | 2.7 |  |  |
| COP9 signalosome | 2.3 |  |  |
| Microtubule | 1.5 |  |  |

To illustrate the wealth of information provided in the dataset, we show examples of proteins from pathways discussed in **Table 1**, specifically those that are iCrMN- or iSpMN-enriched (**Figure 4**). For example, **Figure 4A** shows the expression signatures for several solute carriers that we identified; other solute carriers are shown in **Supplementary Figure S4.** Solute carriers play important roles in both the physiology and degeneration of the nervous system ^24–26^. Slc24a2 is a neuronal sodium/calcium exchanger and important for synaptic plasticity underlying learning and memory ^27^. Slc25a24 is an ATP-Mg/Pi carrier which regulates the adenine nucleotide pool in the mitochondrial matrix and therefore ATP production ^28^. It is important for axon and presynaptic terminal function ^29^. Slc9a1 plays an important role in intracellular pH homeostasis ^30^. Deletion and mutation of Slc9a1 function in mice result in neurodevelopmental disorders and early death^31,32^.

**Figure 4.**
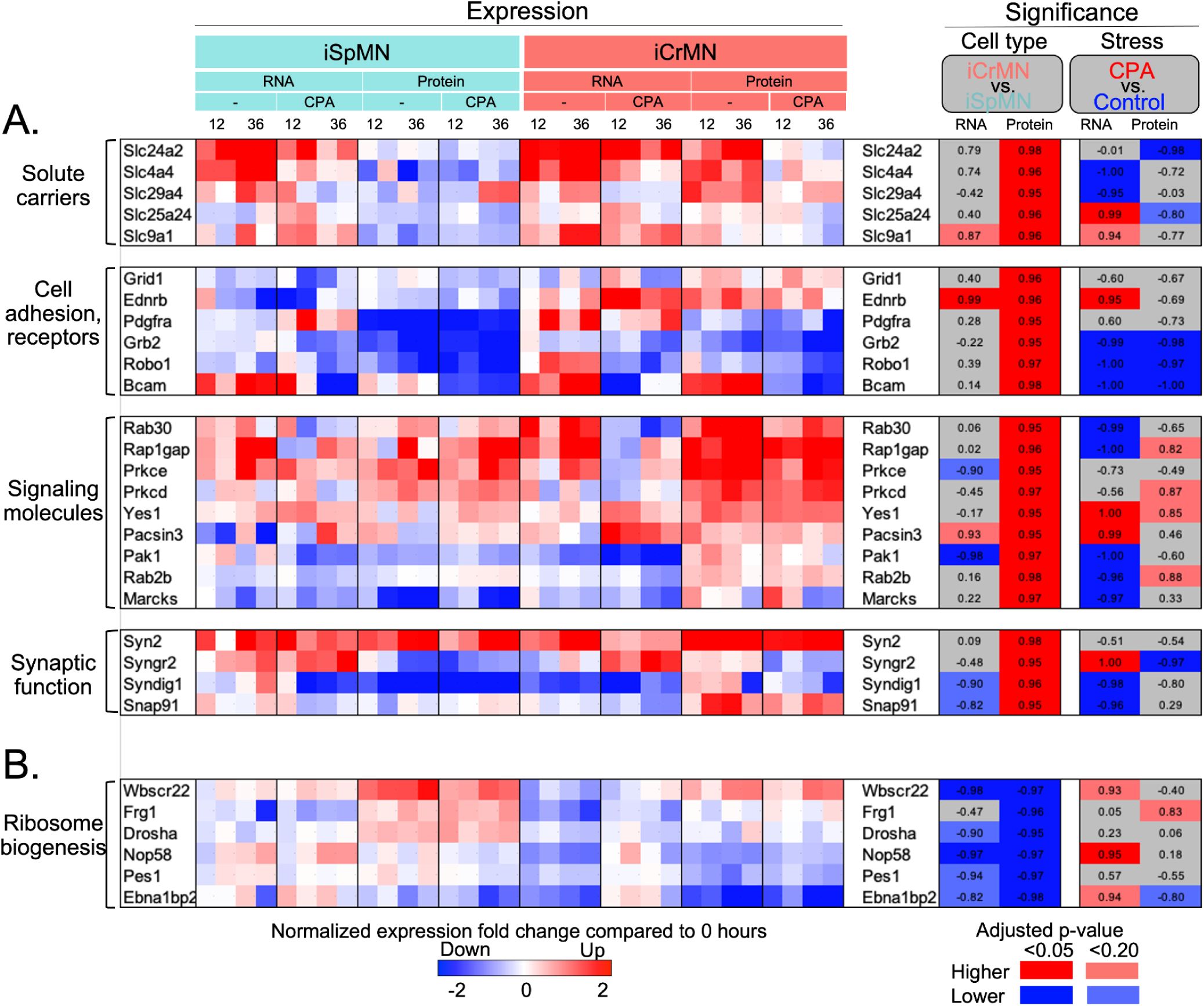
Example proteins from pathways with significant cell type enrichment. Heatmaps showing the expression (left) and significance values (right) for example proteins chosen from pathways enriched in genes with specific expression signatures. Expression differences (left) are Z-score normalized log-transformed ratios of the expression value at 12 or 36 hours compared to 0 hours of treatment with cyclopiazonic acid or DMSO. Significance values (right) are directed adjusted p-values: shades of red if the direction of the expression difference is positive (enriched in iCrMN), shades of blue if the direction is negative (enriched in iSpMN). Adjusted p-values were also thresholded at 0.05 and 0.20 as shown in the legend. Grey indicates no significant difference. Examples were chosen from pathways enriched as described in **Table 1** and indicated by red and blue boxes. **A.** Example genes with significantly higher protein levels in iCrMNs compared to iSpMNs (adjusted p-value < 0.05). **B.** Example genes with significantly higher protein levels in iSpMNs compared to iCrMNs (adjusted p-value < 0.05). CPA - cyclopiazonic acid; iSpMN - induced spinal motor neurons; iCrMN - induced cranial motor neurons

**Figure 4A** also shows examples of cell adhesion molecules enriched in iCrMNs compared to iSpMNs. Cell adhesion molecules play important roles in axon migration and the dynamics of cells in the nervous system ^33–35^. Bcam is a membrane glycoprotein that mediates interaction between cell membranes with laminin α-5 chain ^36,37^. As the laminin α-5 chain is enriched in the microenvironment of the nervous system, Bcam supports general neuronal functioning. Cntn2 is a neuron-specific glycoprotein and plays important roles in neurodevelopment, neuronal migration, and axon fasciculation ^38^. In humans, *CNTN2* mutation is correlated with familial cortical myoclonic tremor and epilepsy ^39^.

**Figure 4A** further highlights other receptors. Robo1 is a member of the family of Robo transmembrane receptors and also a cell adhesion molecule. Robo1 receptors induce actin cytoskeletal rearrangements in projecting axons ^40^, regulate axon fasciculation, and participate in neuron migration ^41^. Grid1 is involved in the formation of excitatory synapses in the cerebellum and hippocampus by activating Ras signaling ^42,43^ and in pruning of the hippocampus and medial prefrontal cortex ^44^. Ednrb is a G protein coupled receptor for endothelins which regulate development of the vertebrate-specific neural crest and its derivatives ^45,46^.

Finally, **Figure 4A** shows examples of signaling molecules, many of which are membrane associated, i.e. consistent with the enrichment described in **Table 1**. Marcks is a scaffold protein which influences synaptic signaling through formation of both pre- and post-synaptic vesicles ^47,48^. Rap1gap is a negative regulator of Rap1/Erk pathway whose overactivation is related to neuron degeneration ^49,50^. Yes1 is a tyrosine kinase associated with focal adhesions and vesicle release ^51^; its specific role in neurons is not known. Grb2 is a scaffold protein that negatively controls signal transduction ^52^. Rab30 and Rab2b are Rab small GTPases and crucial regulators of membrane trafficking and organelle integrity ^53,54^. Both Rab30 and Rab2b regulate the morphology of the Golgi apparatus. Finally, **Figure 4A** shows the Prkce and Prkcd kinase which are highly expressed in brain tissue and are upstream of several neuronal signaling pathways ^55–57^.

Other membrane associated proteins enriched in iCrMNs include components of the lysosomal v-type ATPase (**Supplementary Figure S5**). The lysosomal v-type ATPase regulates lysosomal pH and is involved in various neuron cell functions ^58^. Its dysfunction is part of several disorders, including neurodegeneration^58^.

**Figure 4B** shows examples of proteins which are significantly iSpMN-enriched (adjusted p-value < 0.05). The genes are annotated as functioning in ribosome biogenesis and general RNA processing. **Supplementary Figure S6** shows an expanded list of genes annotated as from these pathways. Notably, the iSpMN-enrichment does not apply to ribosome subunits themselves (**Supplementary Figures S7, S8**). RNA processing is critical for neurological physiology, and the impairment of this process is detrimental for nervous cell function ^59,60^. Wbscr22 promotes the processing of nuclear rRNA precursors and the nuclear export of pre-40S subunits ^61,62^. Frg1 is involved in RNA biogenesis, mRNA transport, and cytoplasmic mRNA localization ^63^. Nop58 is a core component of the snoRNP complex which catalyzes post-transcriptional modifications in ribosomal RNAs (rRNAs)^64,65^. Pes1 controls the nucleolar localization and maturation of the 60S ribosomal subunit ^66,67^. Ebna1bp2 works as a scaffold and is required for pre-rRNA processing and for ribosomal subunit assembly ^68,69^.

Drosha is a class 2 ribonuclease III known to regulate miRNA precursor processing ^70^ and another example of significantly iSpMN-enriched proteins (adjusted p-value < 0.05, **Figure 4B**). miRNA-based post-transcriptional regulation is involved in neuronal differentiation and physiology ^71^. Drosha also has non-canonical roles in promoting post-transcriptional regulation of noncoding and coding RNAs ^72^ and in ribosome biogenesis ^73^. Further, Drosha can recognize and process mRNAs whose sequence and structure are similar to those of pri-miRNAs ^72^. The mRNAs are then cleaved by the miRNA machinery. One example for this mode of regulation is Neurogenin 2 mRNA, which is a key transcription factor for neuron differentiation ^74^. Therefore, Drosha might affect neuronal cell differentiation and physiology via both canonical and non-canonical pathways.

**Table 1** also describes functions enriched amongst RNAs and proteins that changed significantly in response to stress, regardless of the cell type. As expected, many transcripts and proteins significantly induced in response to the treatment are strongly enriched for proteins functioning in the UPR, the response to accumulation of misfolded proteins in the endoplasmic reticulum. In comparison, mRNAs and proteins reduced in response to stress had functions related to the proper working of neuronal cells: they were membrane proteins, including cell adhesion molecules and ion channels. Transcripts from cholesterol biosynthesis, the citric acid cycle, COP9 signaling, and microtubule formation were also significantly lowered in response to stress (FDR<0.05, **Table 1**); **Supplementary Figures S9** and **S10** show examples of genes from these functions. Strikingly, genes from the citric acid cycle and oxidative phosphorylation decrease at the RNA level in response to stress, but not at the protein level (**Supplementary Figure S9**) - and our earlier work hypothesizes on possible translation regulation of these genes ^20^.

### Cell type specific proteasome functionality

Next, we explored an intriguing observation in our data: proteins of the 20S core proteasome particle were significantly enriched in iCrMNs compared to iSpMNs, while this was not the case for proteins of the regulatory particle (**Table 1**). The proteolytic activity of the 20S core particle is controlled by regulatory complexes, such as the 19S or PA28/PA200 particles ^75^. Capping of one or both ends of the 20S particle by the 19S particle forms a new stable complex, called the 26S proteasome which recognizes and degrades ubiquitinated proteins. The 26S particle is therefore responsible for targeted protein degradation. Alternatively, the 20S particle can also be activated by a transient interaction with the PA28 particle. The activated 20S particle can then recognize and degrade peptides and damaged proteins without the need for their ubiquitination ^75^. Therefore, the observed higher abundance of proteins of the 20S core particle was puzzling, in particular as our earlier work had identified higher levels of ubiquitinated proteins in iCrMNs upon proteasome inhibition ^8^ which would require the 26S particle for their degradation.

To investigate these findings further, we first inspected the expression patterns of the genes in more detail (**Figure 5A**). Indeed, while many transcripts encoding members of the 20S and 19S particles were down-regulated in iCrMNs compared to iSpMNs, only proteins from the 20S core particle, but not from the 19S or PA28 particles, showed significantly increased levels in iCrMNs. Proteins from the 19S and PA28 particles were induced under stress, while this was not the case for proteins from the core particles. These results were supported by western blots analysis of individual proteins (**Supplementary Figure S11**). We did not observe similarly consistent expression patterns for genes involved in the lysosome/autosome system (**Supplementary Figure S12**).

**Figure 5.**
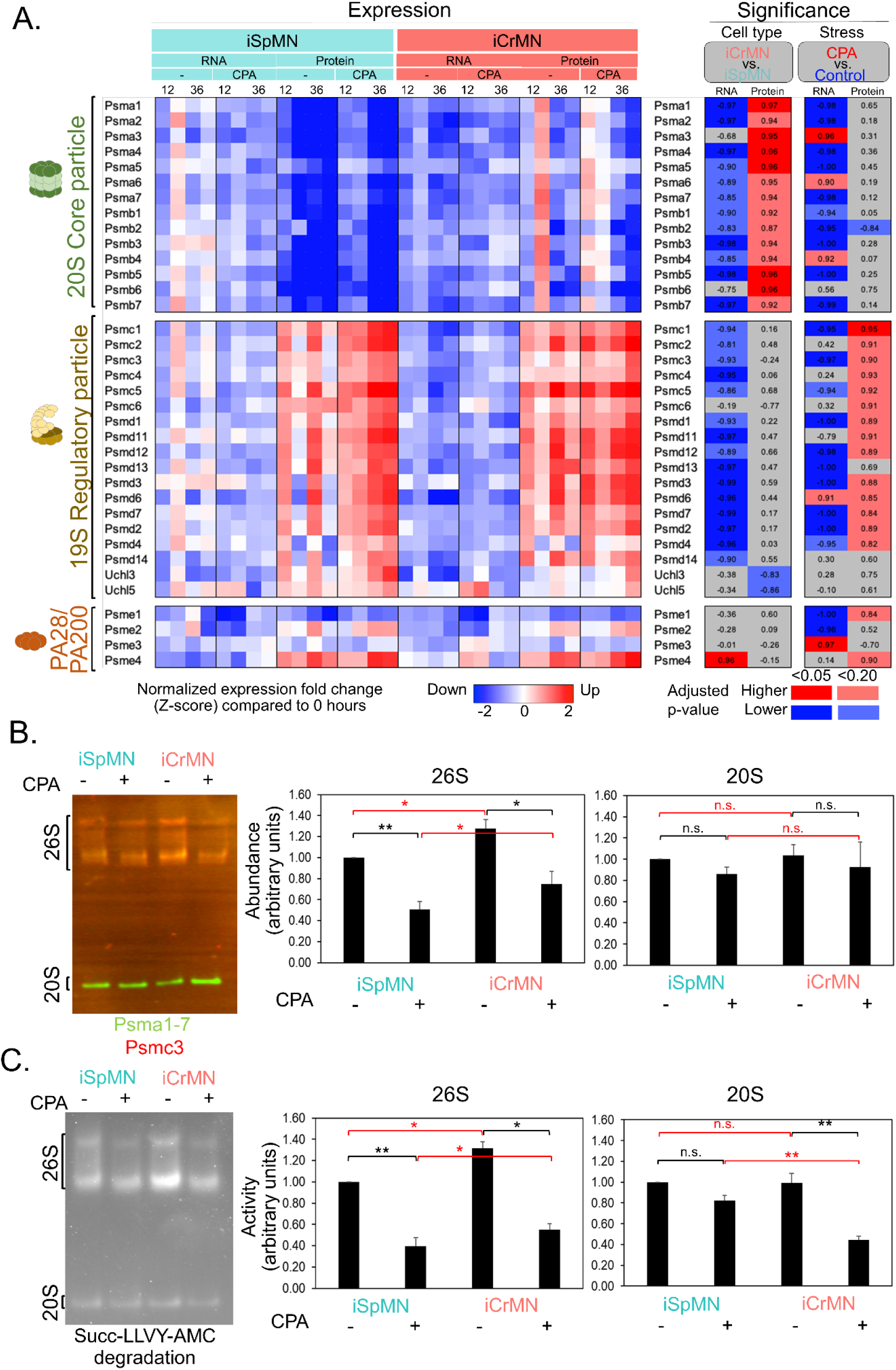
Abundance and activity of proteasome subunits and particles. **A.** Heatmaps showing the expression (left) and significance values (right) for subunits from the 20S core, and the 19S and PA28/PA200 regulatory particles. Expression differences (left) are Z-score normalized log-transformed ratios of the expression value at 12 or 36 hours compared to 0 hours of treatment with cyclopiazonic acid or DMSO. Significance values (right) are directed adjusted p-values: shades of red if the direction of the expression difference is positive (enriched in iCrMN), shades of blue if the direction is negative (enriched in iSpMN). Adjusted p-values were also thresholded at 0.05 and 0.20 as shown in the legend. Grey indicates no significant difference. **B.** Immunoblot of native protein extracts from both iSpMN and iCrMN cells treated with DMSO or CPA. Proteins had been gel-separated under non-denaturing conditions, transferred to a membrane to quantify the abundance of different proteasome particles using specific antibodies against: Psmc3 (red) for 26S particles and Psma1-7 (green) for 20S. **C.** Proteasome in-gel activity assay of native protein extracts from both iSpMN and iCrMN cells treated with DMSO or CPA. Proteins had been gel-separated under non-denaturing conditions prior to measuring the activity of the different proteasome particles by the turnover of the substrate peptide Suc-LLVY-Amc. The graphs next to blots in **B.** and **C.** show the quantitative results. Intensities measured in the gels were normalized by calculating the ratio between the value measured for individual bands and the sum of all measurements taken for the gel (membrane). All graphs show normalized intensities in relationship to the iSpMN control sample, displayed as the mean and standard error from n=5. Red brackets indicate differences between cell types; black brackets indicate changes in response to stress. * - p-value < 0.05; ** - p-value < 0.01; CPA - cyclopiazonic acid; iSpMN - induced spinal motor neurons; iCrMN - induced cranial motor neurons; n.s. - not significant

Therefore, we proceeded to characterize the proteasome further, i.e. with respect to the differential abundance of the assembled 20S and 26S particles. To do so, we separated cell extract under native conditions, i.e. keeping protein complexes intact, and then probed for the presence of individual proteins of the 20S and 19S particles. We observed the 20S and 26S at the expected molecular weight of ∼750 kDa and ∼2000-3000 kDa, respectively, indicating successful isolation of the intact complexes (**Figure 5B**).

Upon quantitation of the abundances, we found that CrMNs had significantly more fully assembled 26S proteasome than iSpMNs, but similar levels of 20S proteasome (p-value < 0.05, **Figure 5B**). Stress reduced the abundance of both the 26S and 20S particles in both cell types, even if not significantly for the 20S in iSpMN. As a consequence, iCrMNs maintained higher 26S levels compared to iSpMNs, but similar or lower levels of 20S particles, under both normal and stress conditions (red brackets in **Figure 5B**). The differential abundance of proteins of the 20S particle observed in **Figure 5A** is therefore explained by the fact that these proteins contribute to both the 26S and 20S particles.

As the higher abundance of 26S particles in iCrMNs suggested also higher proteolytic activity, we next investigated proteasome activity in detail. To do so, we first assayed overall proteasome activity using a conventional plate assay and fluorescently labeled substrates. We found an increased proteasome activity in iCrMNs compared to iSpMNs (**Supplementary Figure S13**), as reported previously ^8^. Intriguingly, stress reduced proteasome activity in both cell types by ∼50% - expanding prior results.

Next, we developed an in-gel assay which probed the differently sized proteasome particles for their proteolytic activity (**Figure 5C**)^76^. In contrast to previous work, this assay quantitates particle-specific activities. For both the 26S and 20S particles and for both cell types, proteolytic activity decreased in response to stress. Similar to what we had observed for 26S abundance, iCrMNs showed significantly higher 26S activity under both normal and stress conditions (p-value < 0.05), indicating the dominant use of ubiquitin-dependent degradation (red brackets in **Figure 5C**). In comparison, iSpMNs showed significantly higher activity of the 20S particle under stress conditions (p-value < 0.05), indicating that they rely more on ubiquitin-independent degradation. The observation expands previous observations to demonstrate that the superior ability of iCrMN to remove damaged proteins relies on heightened abundance and activity of the fully assembled 26S proteasome.

### Regulators of proteasome biogenesis, assembly, and activity

The abundance and activity of the 26S and 20S proteasome particles are finely controlled by several regulators of proteasome biogenesis, assembly, and activity. Alterations, e.g. in assembly chaperones, result in strong defects ^77^. Therefore we investigated the potential role of proteasome regulators in establishing the surplus of 26S particles in iCrMNs, focussing on two classes: chaperones of proteasome biogenesis (**Figure 6A**) and activity regulators (**Figure 6B**). Examples of other proteasome regulators are shown in **Supplementary Figure S14**.

**Figure 6.**
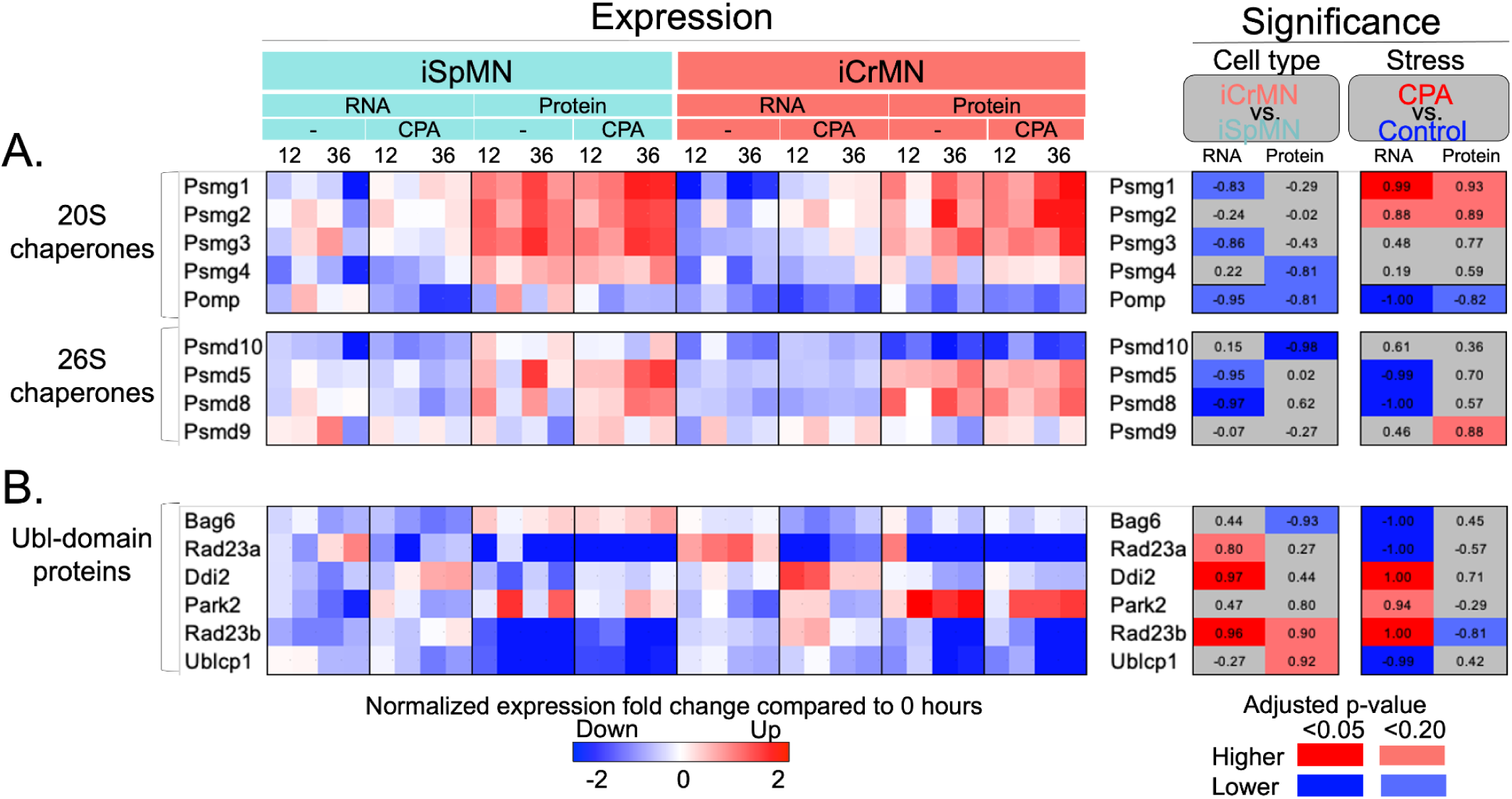
Expression signatures of regulators of proteasome biogenesis, assembly, and activity. Heatmaps showing the expression (left) and significance values (right) for example proteasome regulators, i.e. chaperones (**A.**) and Ubl-domain containing proteins (**B.**). Expression differences (left) are Z-score normalized log-transformed ratios of the expression value at 12 or 36 hours compared to 0 hours of treatment with cyclopiazonic acid or DMSO. Significance values (right) are directed adjusted p-values: shades of red if the direction of the expression difference is positive (enriched in iCrMN), shades of blue if the direction is negative (enriched in iSpMN). Adjusted p-values were also thresholded at 0.05 and 0.20 as shown in the legend. Grey indicates no significant difference. * - p-value < 0.05; ** - p-value < 0.01; CPA - cyclopiazonic acid; iSpMN - induced spinal motor neurons; iCrMN - induced cranial motor neurons; n.s. - not significant

The 20S core proteasome requires five chaperones for its full assembly ^78,79^, shown in **Figure 6A**. Psmg1 to Psmg4 show similar protein expression patterns, without significant differences between cell types. Psmg4 (Pac4) is a 20S proteasome assembly chaperone which forms homodimers or heterodimers with Psmg3 ^78,79^. Pomp facilitates 20S maturation at the ER ^80,81^. Psmg1, Psmg2, and Pomp showed a mild stress response (p-value < 0.20).

**Figure 6A** also shows four chaperones of the 26S particle ^82–87^: Psmd5 and Psmd8 (Paaf1) to Psmd10. The chaperones bind to proteins of the Psmc family which are part of the 19S regulatory particle ^88,89^). Only Psmd10 showed a significant difference between the two cell types in our data: it was iSpMN-enriched (adjusted p-value < 0.05).

**Figure 6B** shows a selection of proteasome activity regulators, specifically Ubiquitin-like (UbL)-domain proteins as examined in ref. ^90^: Bag6, Rad23a, Rad23b, Ddi2, Park2, and Ublcp1. UbL-domain proteins represent a conserved group of diverse proteins that share a common property with respect to interactions with the 26S proteasome and subsequent increase in its activity ^90–92^. Bag6 was the only protein with mildly higher levels in iSpMNs, while Rad23b and Ublcp1 both had higher levels in iCrMNs (adjusted p-value < 0.20).

### The role of the nuclear proteasome regulator Ublcp1

Next, we investigated Ublcp1 in more detail, as it is a little-characterized proteasome regulator but was differentially abundant in the two cell types (**Figure 6B**). Western blotting against Ublcp1 confirmed observations made by mass spectrometry: iCrMNs had higher levels of the protein than iSpMNs (**Figure 7A**). We then tested Ublcp1’s general impact on cellular survival, using a Ublcp1 inhibitor (**Figure 7B**). We found that iCrMNs were more sensitive to Ublcp1 inhibition than iSpMNs under both normal and stress conditions: Ublcp1 inhibition led to significantly less survival in iCrMNs (p-value < 0.05), but not in iSpMNs (purple brackets in **Figure 7B**).

**Figure 7.**
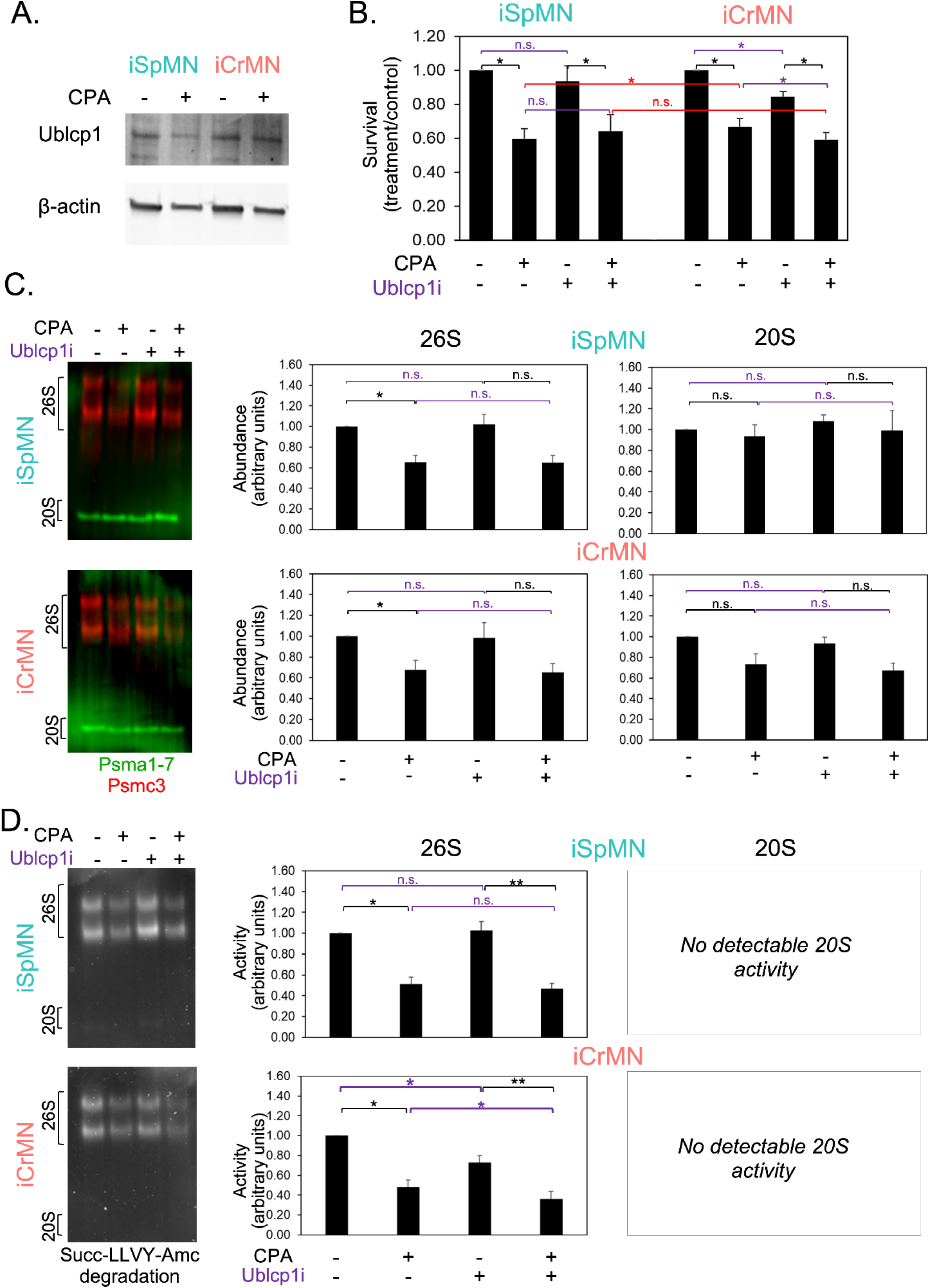
Ublcp1 as a contributor to the differential stress resistance of motor neurons. **A.** Immunoblot for the detection of Ublcp1 levels in extracts from iSpMNs and iCrMNs treated with DMSO or CPA for 12 hours. β-actin was used as loading control. **B.** Survival of iSpMNs and iCrMNs maintained without or with CPA and with or without the Ublcp1 inhibitor. The survival was measured after 48 hours of treatment using Alamar Blue reduction. Survival is plotted in relationship to controls (-/-), as the mean and standard error from n=5. Black brackets refer to stress differences, purple brackets to differences with respect to Ublcp1 inhibition, and red brackets to cell type differences. **C/D**. Nuclear proteasome abundance and activity of proteins from both iSpMN and iCrMN cells treated with DMSO or CPA. Protein samples had been separated in SDS-free gels under non-denaturing conditions prior to measuring the abundance with antibodies against Psmc3 (red) for 26S/30S particles and against Psma1-7 (green) for 20S (**C.**), and activity of different particles by the turnover of the substrate peptide Suc-LLVY-Amc (**D.**). The graphs next to the blots show the quantitative results for both 20S and 26S abundance and activity. Intensities were normalized by calculating the ratio between the value measured for individual bands and the sum of all measurements taken for the entire gel. All graphs show normalized intensities in relationship to the iSpMN control sample, displayed as the mean and standard error of the mean for n=5. Black brackets refer to stress differences, and purple brackets to differences with respect to Ublcp1 inhibition. * - p-value < 0.05; ** - p-value < 0.01; CPA - cyclopiazonic acid; iSpMN - induced spinal motor neurons; iCrMN - induced cranial motor neurons; n.s. - not significant; Ublcp1i - Ublcp1 inhibitor

Next, we evaluated the specific impact of Ublcp1 on proteasome activity in the two cell types. As Ublcp1 is exclusively localized to the nucleus ^93^, and is thought to affect the abundance of the 26S compared to the 20S particle ^93^, we developed a protocol in which we first fractionated nuclear and cytoplasmic protein complexes, and then used native gel separation as well as in-gel activity assays as described above to estimate activities of the different proteasome particles. **Figure 7C** and **Supplementary Figure S15** show the results for the nuclear and cytoplasmic proteasome particles, respectively. Note that activity of the 20S particle in the nuclear fraction was below the detection limit for both cell types.

Similar to what we had observed for the whole cell extract (**Figure 5**), proteotoxic stress reduced activity of the 26S particle in both cell types in both the cytoplasmic and nuclear fraction (black brackets in **Figure 7C, Supplementary Figure S15**). In comparison, only the activity of the cytoplasmic 20S particle from iCrMNs, but not from iSpMNs, was significantly reduced under stress (p-value < 0.05, **Supplementary Figure S16**).

Consistent with what is known about Ublcp1’s nuclear localization ^93^, we observed no measurable sensitivity of the cytoplasmic proteasome particles to Ublcp1 inhibition (purple brackets in **Supplementary Figure S15**). However, intriguingly, for the nuclear proteasome iSpMNs and iCrMNs showed different sensitivities to Ublcp1 inhibition: 26S proteasome activity was significantly lower under both normal and stress conditions under Ublcp1 inhibition in iCrMNs, but not in iSpMNs (p-value < 0.05, purple brackets in **Figure 7C**). The results suggest that Ublcp1’s function is more important for nuclear 26S proteasome activity in iCrMNs than in iSpMNs, possibly supporting the superior stress resistance of iCrMNs. The higher survival rates of iCrMNs in response to stress vanish when Ublcp1 is inhibited.

## Discussion

We present an in-depth, time-resolved dataset describing transcriptomic and proteomic changes in response to proteotoxic stress in an induced spinal and cranial motor neuron system. We identified hundreds of significant expression differences at the RNA or protein level from a core set of >8,200 genes that characterize the two cell types as well as the stress response (**Figure 1-3**). The results of our analyses represent a resource for the community to generate new hypotheses on the differential sensitivity of iSpMN and iCrMN to misfolded protein stress - which is directly relevant to further our understanding of the molecular basis of ALS resistance of a subset of rostral and ocular motor neurons, compared to ALS-sensitive spinal motor neurons. We observed an overall strong similarity between the expression profiles of the two cell types (**Figure 2**), consistent with their highly similar physiological features ^21^. While transcriptome and proteome remodeling was dominated by the response to proteotoxic stress, we also identified several hundreds of differences between the two cell types (**Figure 1**). These differences largely manifested at either the RNA or protein level, but not at both (**Figure 3**).

We focussed on differences between the two cell types with respect to protein expression levels. Proteins enriched in iCrMNs included many membrane proteins (**Table 1**), for example solute carriers, cell adhesion molecules and receptors, as well as other proteins involved in post- and pre-synaptic vesicle formation and neurotransmitters release (**Figure 4**). The corresponding pathways support overall cellular integrity. For example, neurons rely on solute carriers to fulfill their signaling role ^94,95^: Slc42a2 supports sodium/proton exchange on the plasma membrane, and Slc25a24 controls the efflux of adenine nucleotides ^28^. Both proteins were iCrMN-enriched, but their levels decreased under stress (**Figure 4**), suggesting that their reduction might negatively impact electrolyte and energy balance in the cell ^96,97^.

Other iCrMN-enriched membrane proteins included cell adhesion molecules and membrane receptors (**Table 1**, **Figure 4**) which can affect neuron polarity and the distribution of organelles such as the ER and Golgi apparatus within axonal processes ^98–100^. Membrane localized signaling and synapse related-proteins control neurotransmitter release and synaptic vesicle formation ^47,48,55,57,101,102^ (**Table 1**, **Figure 4**). These functions suggest that several pathways closely related to proper neuronal function are more prevalent in cranial motor neurons. As stress reduces the abundance of some of these proteins, spinal motor neurons might more easily lose these functions and become stress sensitive.

iSpMN-enriched proteins included members of the ribosome biogenesis pathway and RNA processing (**Table 1**, **Figure 4**). Therefore, spinal motor neurons might be more active in protein synthesis and other processes involving RNAs and RNA-protein complexes, such as miRNA-based expression regulation, telomere integrity, or pre-mRNA splicing ^103^. While several links between ribosome biogenesis, RNA processing, and neurodegeneration exist ^104^, the relationship is complex: dysfunctional ribosome biogenesis causes proteotoxic stress ^105^, and several ALS-genes are involved in different aspects of RNA processing and translation ^106^. Reduction in rRNA expression can lead to neuronal cell death ^106^.

Further, it has been shown that elevated levels of Drosha sensitizes the cells to proteotoxic stress ^107^. Conversely, reduced Drosha levels enhance axonal growth and therefore mitigate the effects of Spinal Muscular Atrophy, a neurodegenerative disease similar to ALS ^107^. This role suggests a route by which lower Drosha levels in iCrMNs support their neuronal function.

We explored in more detail the enrichment of subunits of the 20S proteasome core particle in iCrMNs (**Table 1**). This observation was intriguing in the light of the role of proteasomal degradation in proteostasis control, emerging evidence for its role in neuronal survival in ALS ^8,108–111^, and recent findings on the localization of some 20S particles to the neuronal plasma membranes ^112^. Despite extensive testing, we could not find sufficient evidence in support of the presence of a neuronal membrane proteasome ^112^, which could be attributed to the use of a different cell system (*data not shown*).

We identified two main characteristics of the neuronal proteasome network that expanded what is known from literature ^8,108–110^, exploiting highly specific proteasome activity assays. First, we showed that both the activity of the fully assembled 26S particle, but not the 20S core, were higher in induced cranial motor neurons compared to spinal motor neurons (**Figure 5**). Second, we showed that proteotoxic stress reduced the activity of the 26S particle in both cell types (**Figure 5**). This finding is consistent with disassembly of the 26S during oxidative stress which can be a result of the UPR ^113,114^. Therefore, we showed that the 26S particle is the bottleneck in the ability of iCrMNs and iSpMNs to respond to proteostatic stress.

As the 26S, but not the 20S proteasome, rely on ubiquitin-tagging of target proteins ^115^, the results suggest an extensive role of the ubiquitination system in manifesting the differences between iCrMNs and iSpMNs. Indeed, several E3 ubiquitin ligases differed significantly between iCrMNs and iSpMNs (**Supplementary Figure S16**). The most significantly iSpMN-enriched E3 ligase was Fbxo2 (**Supplementary Figure S16**). Fbxo2 recognizes glycoproteins ^116^ and can therefore contribute to the regulation of neuronal function through targeting of neuron glycoproteins for degradation ^117^. A notable example of a significantly iCrMN-enriched E3 ubiquitin ligase was Mid1 (**Supplementary Figure S16**). Mid1 is highly relevant to neurodegenerative diseases ^118,119^ as a major target includes Pp2a, the major Tau phosphatase ^120,121^. As both Pp2a and Tau were also significantly different between iCrMNs and iSpMNs, it is tempting to speculate on a role of the Mid1-Pp2a-Tau relationship in establishing the differences between the two motor neuron types. Indeed, emerging evidence suggests a role of Tau in the ALS pathology ^122^.

Finally, we identified Ublcp1, a little-characterized 26S activator ^93,123,124^, as a new regulator of the differences between iCrMNs and iSpMNs under both normal and stress conditions (**Figure 7**). Importantly, we showed that Ublcp1 impacted the *activity*, but not the *abundance* of the nuclear 26S particle in iCrMNs, and its inhibition abolished the superior survival of iCrMNs compared to iSpMNs under stress (**Figure 7**).

These findings are highly relevant in the light of the role of nuclear proteostasis in maintaining cellular function ^125–127^, in particular in non-dividing cells, such as neurons ^128^. However, few regulators of nuclear proteasome activity and abundance are known, and most studies have been conducted in yeast ^128^. Only very recent work identified a nuclear proteasome import factor, AKIRIN2, which is a highly conserved protein which dimerizes and binds to the 20S particle ^129^. Therefore, our results on Ublcp1 not only suggest a new gene with potential ALS relevance, but also advance our understanding of nuclear proteostasis in general.

We hope that these results inspire many future investigations, as there are many expression signatures whose exploration would go beyond the scope of this analysis. One example is Rad23b which we showed to be both iCrMN-enriched and stress-responsive (**Figure 6A**). Rad23b was first characterized as involved in nucleotide excision repair ^130,131^; however, several studies have now described a role in delivery of ubiquitinated proteins to the proteasome and proteasome activation ^132,133,134^. Therefore, Rad23B might be another factor supporting the iCrMN’s superior stress response.

Other proteasome regulators which might support the differences between iCrMNs and iSpMNs include kinases and phosphatases which control, amongst other functions, the assembly of the 26S particle (**Supplementary Figure S14**). For example, we found that Ampka2 (Prkaa2), Pkg (Prkg1), and Pp2a (Ppp2ca) were iCrMN-enriched. Ampka2 plays key roles in proteasome 26S assembly and activity ^135^, and Pkg-based phosphorylation increases activity of the activation of the cytoplasmic proteasome ^136^. Pp2a counterbalances Pka activity which is a proteasome assembly activator ^137,138^. It is a target of the Mid1 E3 ligase mentioned above. In comparison, the Ask1 (Map3k5) kinase was iSpMN-enriched (**Supplementary Figure S14**). Ask1 mediates proteasomal dysfunction through Psmc3 phosphorylation and its subsequent reduction of its ATPase activity ^139^. The findings on Ask1 are particularly interesting in the light of its emerging role as a therapeutic target for neurodegenerative diseases ^140,141^.

Finally, proteasome function is further modified by interacting proteins, several of which show differential expression in iCrMNs and iSpMNs (**Supplementary Figure S14**). Amongst these, the ubiquitin segregase Valosin-containing protein (Vcp, p97) might perhaps be most interesting (**Supplementary Figure S17**). Vcp is a major component of the ribosome quality control complex (RQC), interacts with the proteasome, and is involved in stress granule formation ^142^, therefore linking three key processes of the response to proteostasis stress. Indeed, we found that the RQC and its trigger complex were stress induced in both cell lines (**Supplementary Figure S17**). Given its role in ALS and related diseases ^143^, investigation of the RQC in the two motor neuron types might be another worthy future undertaking.

In sum, our results represent the first large-scale, unbiased, and quantitative view of the complex transcriptomic and proteomic landscape of two physiologically similar motor neurons responding to proteostasis stress. While we detected many novel differences between the cell types, we showed at the example of the proteasome how these differences are established with respect to an individual pathway. Our results demonstrated that the superior stress resistance of iCrMNs compared to iSpMNs is likely not due to individual genes, but is based on a ‘neuronal proteasome network’ that employs different strategies to ensure a strong response to challenges. Future work will likely expand this network to the entire cell.

## Material and Methods

### Motor neuron differentiation

Mouse embryonic cells (mESC) containing inducible NIP and NIL transcription factor cassettes were cultured as previously described ^8,21^. mESCs were grown in 1% gelatin coated flasks in Advanced DMEM/F12:Neurobasal medium (Thermo Fisher, 12634028, 10888022) supplemented with 2.5% tetracycline-negative fetal bovine serum (vol/vol, Corning 35–075-CV), 1X N2 (Thermo Fisher, 17502–048), 1X B27 (Thermo Fisher, 17504–044), 2 mM L-glutamine (Thermo Fisher, 25030081), 0.1 mM β-mercaptoethanol (Thermo Fisher, 21985–023), 1000 U/ml leukemia inhibitory factor (Sigma Aldrich, ESG1107), 3 μM CHIR99021 (Biovision, 1991) and 1 μM PD0325901 (Sigma, PZ0162-5MG), and maintained in incubators at 37°C, 8% CO2. When cells reached 70% confluence colonies were dissociated in TrypLE (Thermo Fisher, 12605-028) to get single-cell suspension. We seeded ∼500,000 single cells in low binding 100 mm plates (Corning, 08-772-32) in differentiation AK medium ((Advanced DMEM/F12: Neurobasal (1:1) Medium (Thermo Fisher, 12634028, 10888022), 10% Knockout SR (Thermo Fisher, 10828–028), 2 mM L-glutamine (Thermo Fisher, 25030081), and 0.1 mM 2-mercaptoethanol (Thermo Fisher, 21985–023) to induce formation of embryoid bodies (EBs). After 2 days, EBs were transferred to low-attachment 100 mm culture dishes and maintained in AK medium containing 3 μM doxycycline for another 2 days to induce expression of the NIP and NIL transcription factors. The EBs were collected and dissociated in 0.05% Trypsin-EDTA (Thermo Fisher, 25300–120). After dissociation, ∼3 million cells were plated in a 100 mm culture dish (Thermo Fisher, 12-567-650) coated with 0.001% poly-D-lysine (Sigma Aldrich, P0899) and were maintained in motor neuron differentiation medium (Neurobasal medium (Thermo Fisher, 10888022) supplemented with 2% fetal bovine serum (Corning, 35–075-CV), 1X B27 (Thermo Fisher, 17504–044), 0.5 mM L-glutamine (Thermo Fisher, 25030081), 0.01 mM 2-mercaptoethanol, 10 ng/mL GDNF (PeproTech, 450–10), 10 ng/mL BDNF (PeproTech 450–02), 10 ng/mL CNTF (PeproTech, 450–13), 10 μM Forskolin (Fisher, BP2520-5) and 100 μM IBMX (Tocris, 2845). Cells were maintained in incubators at 37°C, 5% CO2.

### Treatment

After 2 days of motor neuron differentiation, we induced proteostatic stress by treating cells with 20 μM cyclopiazonic acid (CPA, Sigma-Aldrich, C1530). Control cells were treated with DMSO. We collected cells at the time points listed for the individual assays. To test for the role of Ublcp1, cells were pretreated for 30 minutes with 2.5 μM Ublcp1 inhibitor (Ublcp1i, EMD Millipore, 5.34380.0001) or DMSO as control for 30 minutes prior to CPA/DMSO treatment as described above.

### Survival assay

Cell survival was determined by Alamar Blue reduction (BioRad, BUF012B) following manufacturer instructions. Cells were seeded in 24 well plates, differentiated and treated as described above. Survival was measured as absorbance at 570 nm and 600 nm in a microplate reader (Tecan, Spark microplate reader) after 4 hours of incubation with the reagent. The percent difference in reduction between treated and control cells was determined following manufacturer instructions. All measurements were normalized relative to the respective control.

### RNA-seq analysis

We collected cells at different three time points post treatments (0, 12, 36 hours). The medium was removed, and the cells were washed with 10 ml chilled PBS. RNA was obtained according to the RNeasy mini kit (Qiagen, 74004) extraction protocol. We also performed in-column DNAse digestion using RNAse-Free DNase (Qiagen, 79254). The entire RNA sample was then used to prepare the RNA-seq library preparation using the RNA HyperPrep Kit (Kapa Biosystem, KK8560) protocol. We used TapeStation (Agilent) to confirm the quality of the samples. We quantified samples using Qubit Fluorometry and sequenced the two replicates using Illumina NextSeq500 in high throughput mode.

After sequencing, we used Picard tools (Illumina Basecalls to Fastq version 2.17.11) and pheniqs (https://github.com/biosails/pheniqs, developed by NYU) for base calling and demultiplexing. The sequenced reads were trimmed to remove the adapter sequences using fastx-toolkit v 0.0.14 (with parameters -t 20 -l 20). Multiple fastq files from the same sample were merged. To remove abundant sequences, the reads were mapped to rDNA, adapter contaminants, PhiX, and polyA/C sequences using bowtie2 2.3.2 ^144^ with default settings. The remaining reads were mapped to GRCm38 - mm10 using tophat v2.1.1. ^145^ with the following parameter settings: --no-coverage-search -G gencode mm10 basic.gtf. The aligned reads were filtered for unique mapping and a length of >25 nucleotides. RNA reads aligning to exons were counted as a measure of mRNA abundance. Read counts were generated using htseq v0.6.1p1. Surrogate variables were estimated and removed via linear modeling (SVA) to remove batch effects ^146^.

### Proteomics analysis

Proteomics samples were processed from the same biological replicates as the samples used for RNA-seq analysis. We collected cells at different time points (0, 12, 36 hours post treatment) by removing the medium and scraping cells off the plate. The cell pellet was washed in an ice–cold PBS buffer. After centrifugation, we resuspended the cell pellets in a lysis buffer containing 0.1% Rapigest (Waters,186002122), in 0.1 M HEPES pH 7 (Sigma Aldrich, H0887), and cOmplete protease inhibitor (Roche, 04693132001). The cells were then sonicated using the Bioraptor system (Diaginode) with protein extraction beads (30sec on/off, 5 cycles) at 4 °C. The protein extract from respective samples was spun down at 13,000 g for 10 mins and supernatant was collected in low adhesion 1.5 mL tubes (Eppendorf). We reduced the sample with 15 mM DTT (Sigma Aldrich, 43816) at 55 °C for 45 min and then alkylated with 55 mM iodoacetamide (Sigma Aldrich, A3221) in the dark at room temperature for 30 min. We digested the samples overnight using sequencing grade modified porcine trypsin (1:50 w/w, Sigma Aldrich, 1700013) on a thermomixer at 37 °C, 200 RPM (Eppendorf). Rapigest surfactant was cleaved by incubating samples with ∼200mM HCl (Sigma-Aldrich, H1758) (30 min, 37 °C). We desalted the digested protein samples on C18 spin tips (Thermo Fisher Scientific, 89870) and dried the peptides under a vacuum.

Peptide concentrations were determined using the Quantitative Fluorometric Peptide Assay (Thermo Fisher Scientific, 23290) and 18 µg peptides per time point were subjected to TMT labeling. TMT 10-plex reagents (Thermo Fisher Scientific, 90110) were dissolved in anhydrous acetonitrile (0.8 mg/ 40 µl) according to the manufacturer’s instructions. Each peptide sample was labeled with 41 µl of the TMT 10-plex label reagent at the final acetonitrile concentration of 30% (v/v). Following incubation at room temperature for 1 hour, the reactions were quenched with 8 µl of 5% hydroxylamine for 15 min. All samples were combined into a new microcentrifuge tube at equal amounts and dried down to remove acetonitrile. TMT-labeled tryptic peptides were subjected to high-pH reversed-phase HPLC fractionation using an Agilent 1200 Infinity Series with a phenomenex Kinetex 5u EVO C18 100A column (100 mm x 2.1 mm, 5 mm particle size). Mobile phase A was 20 mM ammonium formate (Sigma-Aldrich, 70221), and B was 10% ammonium formate (200 mM, Sigma-Aldrich) in 90% acetonitrile (EMD Millipore, AX0156-1), both were adjusted to pH 10. Peptides were resolved using a linear 120 min 0-40% acetonitrile gradient at a 100 μl/min flow rate. Eluting peptides were collected in 2 min fractions. About 70 fractions covering the peptide-rich region were combined to provide 40 samples for analysis. Fractions were combined across the gradient. Combined fractions were dried down using a Concentrator Vacufuge Plus (Eppendorf), desalted with C18 stage-tip, and suspended in 95% MS degree water, 5% acetonitrile, and 0.1% formic acid.

We used an EASY-nLC 1200 (Thermo Fisher Scientific) coupled on-line to a Q Exactive-HF spectrometer (Thermo Fisher Scientific) for LC-MS/MS analysis. Solved A (0.1% formic acid in water) and solved B (80% acetonitrile, 0.5% acetic acid) were used as mobile phases for peptide separation. An EASY Spray 50 cm x 75 µm ID PepMap C18 analytical HPLC column (Thermo Fisher Scientific, ES803) with 2 μm bead size was used for peptide separation in an EASY-nLC 1200 HPLC (Thermo Fisher Scientific). Peptides were separated in a gradient composed as follows: 100 minutes from 2% to 20% solvent B (80% acetonitrile, 0.5% acetic acid), 20 minutes from 20% to 32% solvent B, and 5 minutes from 32% to 95% solvent B. Solvent B was held at 95% for another 10 minutes. High-resolution full MS spectra were acquired at a resolution of 120,000, an Automatic Gain Control (AGC) target of 3e6, with a maximum ion time of 100 ms, and a scan range of 375 to 1500 m/z. Following each full MS scan, 15 data-dependent high-resolution HCD MS/MS spectra were acquired. All MS/MS spectra were collected using a resolution of 60,000, AGC target of 2e5, maximum ion time of 100 ms, one microscan, 1.2 m/z isolation window, fixed first mass of 100 m/z, and NCE of 32.

Raw spectra were processed using Proteome Discoverer 2.2 (Thermo Fisher Scientific) with its integrated search engine Sequest HT. The mouse (GRCm38.p6) sequence file was downloaded from the ENSEMBL database. Sample fractions were grouped by setting in the Experimental Design in Proteome Discoverer. The quantification method used the correction factors provided by the TMT Certificate of Analysis. The mass tolerance of the MS/MS spectra was set to 20 ppm with a posterior global false discovery rate of 1% based on reversed mouse sequences. Additional search parameters included common mass spectrometry contaminants, trypsin specificity for peptide cleavage, up to two missed missed cleavages, and carbamidomethylation of cysteine containing peptides.

TMT quantification was performed at the MS2 level with default parameters, using the reporter ion intensities as estimates for abundance. We applied surrogate variable analysis (SVA) to the entire protein abundance dataset, using the “sva” package in R. The variables representing technical noise and batch effects were removed via linear modeling.

### The core dataset and statistical analysis

We derived a core dataset of normalized transcriptomics and proteomics data for 8,206 genes within complete information across all time points and replicates. Next, we calculated the log base 2 ratios of the measurement at 12 or 36 hours compared to the measurement at time 0. We then normalized across the time course (but for each cell type and RNA and protein separately) by subtracting the average value from each entry and dividing by the standard deviation (Z-score). We visualized and analyzed the data matrix in R studio with the ‘ggplot’ package, using hierarchical clustering, the ‘correlation=Pearson’ and ‘method=Complete’ options. We clustered the core dataset into 25 clusters with the highest coherence in the similarity measure.

We performed 3-way ANOVA for the RNA and protein datasets separately to calculate p-values of differential expression for each gene controlled by three different factors (Cell type, Stress, Time) and their interactions. We then calculated the adjusted p-value (Benjamini-Hochberg correction). Function enrichment of significantly differentially expressed genes (adjusted p-value < 0.05) was conducted with the NIH NCBI DAVID tool box, using a false discovery rate < 0.05 (https://david.ncifcrf.gov/summary.jsp).

### SDS-PAGE and immunoblot

To extract proteins, samples were lysed using a modified RIPA buffer (Sigma Aldrich, R0278). Protein concentration was determined using the DC Protein Assay kit (BioRad, 5000112). We mixed cell lysates with sample buffer (Odyssey, 928-40004) containing 0.2% 2-mercaptoethanol and heated samples 95 °C for 5 minutes. We separated 10 μg protein from each sample in 4-15% Tris-Glycine gels (BioRad, 4561084). We then transferred proteins to a low background PVDF membrane (Millipore, IPFL00010) for 15h at 15V using the TRIS-Glycine buffer (48 mM Tris, 39 mM glycine, 1.3mM SDS with 20% methanol). We then blocked the membranes using the non-mammalian blocking buffer TTBS (LiCOR Biosciences, 927-60001) for 1h at room temperature. Next, the membranes were firs incubated in blocking buffer with primary antibodies against the protein of interest overnight at 4 °C, and then with mouse or rabbit secondary antibodies (LiCOR Biosciences, 925-32210 and 926-68073) in TBS for 1h at room temperature. The membrane images were captured with an Odyssey Clx (LiCOR Bio sciences). Quantification of band densitometry was performed using the ImageJ software. Measurements were normalized with the endogenous control β-actin.

We used the following primary antibodies: Psma1-7 (1:1000, Enzo Life Sciences BML-PW8195-0100), Ublcp1 (1:1000, Thermo Scientific PA5-58859) and β-actin (1:10000, Sigma-Aldrich A5441).

### Proteasome activity assays

To extract total native protein, we first extracted proteins under native conditions, as described recently ^76^. First, total protein was extracted from the cell lysate using lysis buffer (50 mM Tris, 1M DTT, 10 mM NaCl, 5 mM MgCl_2_, 0.1% Igepal, 10% Glycerol). We then determined the total protein concentration with the DC Protein assay kit and added lysis buffer to achieve a final protein concentration of 0.5 μg/μl.

To extract the cytoplasmic and nuclear native fractions, we adapted a protocol using differential detergent lysis of cellular fractions ^147^. Cells were resuspended in the extraction buffer (100 mM PIPES, 100 mM NaCl, 3 mM MgCl_2_, 3 mM EDTA, 0.015% Digitonin). To monitor the degree of cell lysis, we used trypan blue staining and evaluation under the microscope, to follow cell lysis. Once ∼95% of the cells were lysed, we centrifuged the suspension at 480g for 10 minutes at 4 °C, and transferred the supernatant containing the cytoplasmic fraction to a fresh tube for snap freezing. We then washed the pellet 3 times with extraction buffer containing 0.05% Igepal prior to resuspension in nuclear extraction buffer (100 mM PIPES, 100 mM NaCl, 3 mM MgCl_2_, 3 mM EDTA, 0.1% Igepal). The samples were kept on ice for 30 minutes, the extract was sonicated and centrifuged at 6780g for 10 minutes at 4 °C. We then transferred the supernatant containing the nuclear extract to a fresh tube prior to snap freezing. All samples were kept at -80 °C until further use. Prior to further analysis, we determined protein concentrations using the DC Protein assay kit and added lysis buffer to obtain a final protein concentration of 0.5 μg/μl for cytoplasm extracts and 0.2 μg/μl for nuclear extracts.

To measure activity of the 20S and 26S particles in either the total or the cytoplasmic and nuclear extracts, we resuspended 15 μl of the respective protein extracts in loading buffer (0.01% Orange G, 8.7% Glycerol, 50 mM Tris, 2 mM DTT, 2 mM ATP). We then separated protein complexes using 3-8% Tris-acetate gels (Thermo Fisher, EA0378) and native running buffer (90 mM Tris Base, 90mM boric acid, 2 mM EDTA-Na_2_, 415mM ATP, 2 mM MgCl_2_, 0.5 mM DTT). Electrophoretic separation was performed for 4 hours at 4 °C and 150V. We then incubated the gel in reaction buffer (Tris 50mM, 10 mM MgCl_2_ Hydrate, 1mM ATP, 1mM DTT) and 48 μM Suc-LLVY-AMC for 30 minutes at room temperature. We then incubated samples for another 30 minutes at 37 °C in the dark, and read plates at A_360_ex./ A_460_em. Using a Gel Doc (BioRad).

To evaluate the abundance of the 20S and 26S particles in the respective samples, we denatured proteins in the gels after the activity assays by incubation in solubilization buffer (2% SDS, 66 mM Na_2_Co_3_, 1.5% 2-mercaptoethanol) for 10 minutes. Proteins were then transferred to a low background PVDF membrane for 16 hours at 40 mAV in TRIS-glycine buffer (48 mM Tris, 39 mM glycine, 1.3 mM SDS with 20% methanol). We then blocked the membranes using the non-mammalian blocking buffer TTBS (LiCOR Biosciences, 927-60001) for 1h at room temperature. Next, the membranes were first incubated in blocking buffer with primary antibodies against the protein of interest overnight at 4 °C, and then with mouse or rabbit secondary antibodies (LiCOR Biosciences, 925-32210 and 926-68073) in TBS for 1h at room temperature. The membrane images were captured with an Odyssey Clx (LiCOR Bio sciences). Quantification of band densitometry was performed using ImageJ software, and bands were normalized by the sum of the bands intensity within the gel.

We used the following primary antibodies: Psma1-7 (1:1000, Enzo Life Sciences BML-PW8195-0100), Psmc3 (1:1000, Atlas Antibodies HPA006065).

## Data availability

Transcriptomics data has been deposited in NCBI’s Gene Expression Omnibus ^148,149^ with the accession number GSE150397. The mass spectrometry data including the Proteome Discoverer output files have been deposited in the PRIDE ^150^ repository with the identifier PXD018070.

## References

1. Fu, H., Hardy, J. & Duff, K. E. Selective vulnerability in neurodegenerative diseases. Nat. Neurosci. 21, 1350–1358 (2018).

2. Saxena, S. & Caroni, P. Selective neuronal vulnerability in neurodegenerative diseases: from stressor thresholds to degeneration. Neuron 71, 35–48 (2011).

3. Nijssen, J., Comley, L. H. & Hedlund, E. Motor neuron vulnerability and resistance in amyotrophic lateral sclerosis. Acta Neuropathol. 133, 863–885 (2017).

4. Brown, R. H. & Al-Chalabi, A. Amyotrophic Lateral Sclerosis. N. Engl. J. Med. 377, 162–172 (2017).

5. Okamoto, K. et al. Oculomotor nuclear pathology in amyotrophic lateral sclerosis. Acta Neuropathol. 85, 458–462 (1993).

6. Lawyer, T. & Netsky, M. G. Amyotrophic lateral sclerosis: a clinicoanatomic study of fifty-three cases. AMA Arch. Neurol. (1953).

7. Kanning, K. C., Kaplan, A. & Henderson, C. E. Motor neuron diversity in development and disease. Annu. Rev. Neurosci. 33, 409–440 (2010).

8. An, D. et al. Stem cell-derived cranial and spinal motor neurons reveal proteostatic differences between ALS resistant and sensitive motor neurons. Elife 8, (2019).

9. Hetz, C. & Mollereau, B. Disturbance of endoplasmic reticulum proteostasis in neurodegenerative diseases. Nat. Rev. Neurosci. 15, 233–249 (2014).

10. Smith, H. L. & Mallucci, G. R. The unfolded protein response: mechanisms and therapy of neurodegeneration. Brain 139, 2113–2121 (2016).

11. Scheper, W. & Hoozemans, J. J. M. The unfolded protein response in neurodegenerative diseases: a neuropathological perspective. Acta Neuropathol. 130, 315–331 (2015).

12. Soto, C. Unfolding the role of protein misfolding in neurodegenerative diseases. Nat. Rev. Neurosci. 4, 49–60 (2003).

13. Navone, F., Genevini, P. & Borgese, N. Autophagy and Neurodegeneration: Insights from a Cultured Cell Model of ALS. Cells 4, 354–386 (2015).

14. Keskin, I. et al. Effects of Cellular Pathway Disturbances on Misfolded Superoxide Dismutase-1 in Fibroblasts Derived from ALS Patients. PLoS One 11, e0150133 (2016).

15. Montibeller, L. & de Belleroche, J. Amyotrophic lateral sclerosis (ALS) and Alzheimer’s disease (AD) are characterised by differential activation of ER stress pathways: focus on UPR target genes. Cell Stress and Chaperones vol. 23 897–912 Preprint at https://doi.org/10.1007/s12192-018-0897-y (2018).

16. Hardiman, O. et al. Amyotrophic lateral sclerosis. Nat Rev Dis Primers 3, 17071 (2017).

17. Kagias, K., Nehammer, C. & Pocock, R. Neuronal responses to physiological stress. Front. Genet. 3, 222 (2012).

18. Peker, N. & Gozuacik, D. Autophagy as a Cellular Stress Response Mechanism in the Nervous System. J. Mol. Biol. 432, 2560–2588 (2020).

19. Smith, M. I. & Deshmukh, M. Endoplasmic reticulum stress-induced apoptosis requires bax for commitment and Apaf-1 for execution in primary neurons. Cell Death Differ. 14, 1011–1019 (2007).

20. Rendleman, J. et al. New insights into the cellular temporal response to proteostatic stress. Elife 7, (2018).

21. Mazzoni, E. O. et al. Synergistic binding of transcription factors to cell-specific enhancers programs motor neuron identity. Nat. Neurosci. 16, 1219–1227 (2013).

22. Moro, S. G., Hermans, C., Ruiz-Orera, J. & Mar Albà, M. Impact of uORFs in mediating regulation of translation in stress conditions. Preprint at https://doi.org/10.21203/rs.3.rs-199549/v1.

23. Young, S. K., Willy, J. A., Wu, C., Sachs, M. S. & Wek, R. C. Ribosome Reinitiation Directs Gene-specific Translation and Regulates the Integrated Stress Response. J. Biol. Chem. 290, 28257–28271 (2015).

24. Ayka, A. & Şehirli, A. Ö. The Role of the SLC Transporters Protein in the Neurodegenerative Disorders. Clinical Psychopharmacology and Neuroscience vol. 18 174–187 Preprint at https://doi.org/10.9758/cpn.2020.18.2.174 (2020).

25. Brzica, H., Abdullahi, W., Ibbotson, K. & Ronaldson, P. T. Role of Transporters in Central Nervous System Drug Delivery and Blood-Brain Barrier Protection: Relevance to Treatment of Stroke. J. Cent. Nerv. Syst. Dis. 9, 1179573517693802 (2017).

26. Nguyen, Y. T. K., Ha, H. T. T., Nguyen, T. H. & Nguyen, L. N. The role of SLC transporters for brain health and disease. Cell. Mol. Life Sci. 79, 20 (2021).

27. Hassan, M. T. & Lytton, J. Potassium-dependent sodium-calcium exchanger (NCKX) isoforms and neuronal function. Cell Calcium 86, 102135 (2020).

28. Harborne, S. P. D., King, M. S., Crichton, P. G. & Kunji, E. R. S. Calcium regulation of the human mitochondrial ATP-Mg/Pi carrier SLC25A24 uses a locking pin mechanism. Sci. Rep. 7, 45383 (2017).

29. Trigo, D., Avelar, C., Fernandes, M., Sá, J. & da Cruz E Silva, O. Mitochondria, energy, and metabolism in neuronal health and disease. FEBS Lett. 596, 1095–1110 (2022).

30. Iwama, K. et al. A novel SLC9A1 mutation causes cerebellar ataxia. J. Hum. Genet. 63, 1049–1054 (2018).

31. Bell, S. M. et al. Targeted disruption of the murine Nhe1 locus induces ataxia, growth retardation, and seizures. Am. J. Physiol. 276, C788–95 (1999).

32. Cox, G. A. et al. Sodium/hydrogen exchanger gene defect in slow-wave epilepsy mutant mice. Cell 91, 139–148 (1997).

33. Mohebiany, A. N., Harroch, S. & Bouyain, S. New insights into the roles of the contactin cell adhesion molecules in neural development. Adv Neurobiol 8, 165–194 (2014).

34. Yamagata, M. & Sanes, J. R. Expanding the Ig superfamily code for laminar specificity in retina: expression and role of contactins. J. Neurosci. 32, 14402–14414 (2012).

35. Borrell, V. et al. Slit/Robo signaling modulates the proliferation of central nervous system progenitors. Neuron 76, 338–352 (2012).

36. Luo, S. et al. Recessive LAMA5 Variants Associated With Partial Epilepsy and Spasms in Infancy. Front. Mol. Neurosci. 15, 825390 (2022).

37. Bartolini, A. et al. BCAM and LAMA5 Mediate the Recognition between Tumor Cells and the Endothelium in the Metastatic Spreading of KRAS-Mutant Colorectal Cancer. Clin. Cancer Res. 22, 4923–4933 (2016).

38. Gurung, S. et al. Distinct roles for the cell adhesion molecule Contactin2 in the development and function of neural circuits in zebrafish. Mech. Dev. 152, 1–12 (2018).

39. Stogmann, E. et al. Autosomal recessive cortical myoclonic tremor and epilepsy: association with a mutation in the potassium channel associated gene CNTN2. Brain 136, 1155–1160 (2013).

40. Lowery, L. A. & Van Vactor, D. The trip of the tip: understanding the growth cone machinery. Nat. Rev. Mol. Cell Biol. 10, 332–343 (2009).

41. Hernández-Miranda, L. R. et al. Robo1 regulates semaphorin signaling to guide the migration of cortical interneurons through the ventral forebrain. J. Neurosci. 31, 6174–6187 (2011).

42. Konno, K. et al. Enriched Expression of GluD1 in Higher Brain Regions and Its Involvement in Parallel Fiber-Interneuron Synapse Formation in the Cerebellum. Journal of Neuroscience vol. 34 7412–7424 Preprint at https://doi.org/10.1523/jneurosci.0628-14.2014 (2014).

43. Tao, W., Díaz-Alonso, J., Sheng, N. & Nicoll, R. A. Postsynaptic δ1 glutamate receptor assembles and maintains hippocampal synapses via Cbln2 and neurexin. Proc. Natl. Acad. Sci. U. S. A. 115, E5373–E5381 (2018).

44. Gupta, S. C. et al. Essential role of GluD1 in dendritic spine development and GluN2B to GluN2A NMDAR subunit switch in the cortex and hippocampus reveals ability of GluN2B inhibition in correcting hyperconnectivity. Neuropharmacology vol. 93 274–284 Preprint at https://doi.org/10.1016/j.neuropharm.2015.02.013 (2015).

45. Bondurand, N., Dufour, S. & Pingault, V. News from the endothelin-3/EDNRB signaling pathway: Role during enteric nervous system development and involvement in neural crest-associated disorders. Dev. Biol. 444 Suppl 1, S156–S169 (2018).

46. Rosanò, L., Spinella, F. & Bagnato, A. Endothelin 1 in cancer: biological implications and therapeutic opportunities. Nat. Rev. Cancer 13, 637–651 (2013).

47. Brudvig, J. J. & Weimer, J. M. X MARCKS the spot: myristoylated alanine-rich C kinase substrate in neuronal function and disease. Front. Cell. Neurosci. 9, 407 (2015).

48. Xu, X.-H. et al. MARCKS regulates membrane targeting of Rab10 vesicles to promote axon development. Cell Res. 24, 576–594 (2014).

49. Paratcha, G. & Ledda, F. The GTPase-activating protein Rap1GAP: a new player to modulate Ret signaling. Cell Res. 21, 217–219 (2011).

50. Subramaniam, S. et al. ERK activation promotes neuronal degeneration predominantly through plasma membrane damage and independently of caspase-3. J. Cell Biol. 165, 357–369 (2004).

51. Kalia, L. V., Gingrich, J. R. & Salter, M. W. Src in synaptic transmission and plasticity. Oncogene 23, 8007–8016 (2004).

52. Ahmed, Z. et al. Grb2 controls phosphorylation of FGFR2 by inhibiting receptor kinase and Shp2 phosphatase activity. J. Cell Biol. 200, 493–504 (2013).

53. Aizawa, M. & Fukuda, M. Small GTPase Rab2B and Its Specific Binding Protein Golgi-associated Rab2B Interactor-like 4 (GARI-L4) Regulate Golgi Morphology. J. Biol. Chem. 290, 22250–22261 (2015).

54. Kelly, E. E. et al. Rab30 is required for the morphological integrity of the Golgi apparatus. Biol. Cell 104, 84–101 (2012).

55. Shirai, Y., Adachi, N. & Saito, N. Protein kinase Cepsilon: function in neurons. FEBS J. 275, 3988–3994 (2008).

56. Zeidman, R., Trollér, U., Raghunath, A., Påhlman, S. & Larsson, C. Protein Kinase Cε Actin-binding Site Is Important for Neurite Outgrowth during Neuronal Differentiation. MBoC 13, 12–24 (2002).

57. Ling, M., Trollér, U., Zeidman, R., Lundberg, C. & Larsson, C. Induction of neurites by the regulatory domains of PKCδ and ε is counteracted by PKC catalytic activity and by the RhoA pathway. Exp. Cell Res. 292, 135–150 (2004).

58. Song, Q., Meng, B., Xu, H. & Mao, Z. The emerging roles of vacuolar-type ATPase-dependent Lysosomal acidification in neurodegenerative diseases. Transl. Neurodegener. 9, 1–14 (2020).

59. Andreassi, C., Crerar, H. & Riccio, A. Post-transcriptional Processing of mRNA in Neurons: The Vestiges of the RNA World Drive Transcriptome Diversity. Front. Mol. Neurosci. 11, 304 (2018).

60. Fusco, C. M. et al. Neuronal ribosomes exhibit dynamic and context-dependent exchange of ribosomal proteins. Nat. Commun. 12, 6127 (2021).

61. Haag, S., Kretschmer, J. & Bohnsack, M. T. WBSCR22/Merm1 is required for late nuclear pre-ribosomal RNA processing and mediates N7-methylation of G1639 in human 18S rRNA. RNA 21, 180–187 (2015).

62. Zorbas, C. et al. The human 18S rRNA base methyltransferases DIMT1L and WBSCR22-TRMT112 but not rRNA modification are required for ribosome biogenesis. Mol. Biol. Cell 26, 2080–2095 (2015).

63. Sun, C.-Y. J. et al. Facioscapulohumeral muscular dystrophy region gene 1 is a dynamic RNA-associated and actin-bundling protein. J. Mol. Biol. 411, 397–416 (2011).

64. Bachellerie, J. P., Cavaillé, J. & Hüttenhofer, A. The expanding snoRNA world. Biochimie 84, 775–790 (2002).

65. Lafontaine, D. L. J. Noncoding RNAs in eukaryotic ribosome biogenesis and function. Nat. Struct. Mol. Biol. 22, 11–19 (2015).

66. Lerch-Gaggl, A. et al. Pescadillo is essential for nucleolar assembly, ribosome biogenesis, and mammalian cell proliferation. J. Biol. Chem. 277, 45347–45355 (2002).

67. Rohrmoser, M. et al. Interdependence of Pes1, Bop1, and WDR12 controls nucleolar localization and assembly of the PeBoW complex required for maturation of the 60S ribosomal subunit. Mol. Cell. Biol. 27, 3682–3694 (2007).

68. Tsujii, R. et al. Ebp2p, yeast homologue of a human protein that interacts with Epstein-Barr virus nuclear antigen 1, is required for pre-rRNA processing and ribosomal subunit assembly. Genes Cells 5, 543–553 (2000).

69. Hirano, Y. et al. Proteomic and targeted analytical identification of BXDC1 and EBNA1BP2 as dynamic scaffold proteins in the nucleolus. Genes Cells 14, 155–166 (2009).

70. Han, J. et al. The Drosha-DGCR8 complex in primary microRNA processing. Genes Dev. 18, 3016–3027 (2004).

71. Thomas, K. T., Gross, C. & Bassell, G. J. microRNAs Sculpt Neuronal Communication in a Tight Balance That Is Lost in Neurological Disease. Front. Mol. Neurosci. 11, 455 (2018).

72. Burger, K. & Gullerova, M. Swiss army knives: non-canonical functions of nuclear Drosha and Dicer. Nat. Rev. Mol. Cell Biol. 16, 417–430 (2015).

73. Liang, X.-H. & Crooke, S. T. Depletion of key protein components of the RISC pathway impairs pre-ribosomal RNA processing. Nucleic Acids Res. 39, 4875–4889 (2011).

74. Knuckles, P. et al. Drosha regulates neurogenesis by controlling neurogenin 2 expression independent of microRNAs. Nat. Neurosci. 15, 962–969 (2012).

75. Tanaka, K. The proteasome: overview of structure and functions. Proc. Jpn. Acad. Ser. B Phys. Biol. Sci. 85, 12–36 (2009).

76. Yazgili, A. S. et al. In-gel proteasome assay to determine the activity, amount, and composition of proteasome complexes from mammalian cells or tissues. STAR Protoc 2, 100526 (2021).

77. Schnell, H. M. et al. Structures of chaperone-associated assembly intermediates reveal coordinated mechanisms of proteasome biogenesis. Nat. Struct. Mol. Biol. 28, 418–425 (2021).

78. Kurimoto, E. et al. Crystal structure of human proteasome assembly chaperone PAC4 involved in proteasome formation. Protein Sci. 26, 1080–1085 (2017).

79. Satoh, T. et al. Molecular and Structural Basis of the Proteasome α Subunit Assembly Mechanism Mediated by the Proteasome-Assembling Chaperone PAC3-PAC4 Heterodimer. Int. J. Mol. Sci. 20, (2019).

80. Fricke, B., Heink, S., Steffen, J., Kloetzel, P.-M. & Krüger, E. The proteasome maturation protein POMP facilitates major steps of 20S proteasome formation at the endoplasmic reticulum. EMBO Rep. 8, 1170–1175 (2007).

81. Witt, E. et al. Characterisation of the newly identified human Ump1 homologue POMP and analysis of LMP7(beta 5i) incorporation into 20 S proteasomes. J. Mol. Biol. 301, 1–9 (2000).

82. Zwickl, P. & Baumeister, W. The Proteasome — Ubiquitin Protein Degradation Pathway. (Springer Science & Business Media, 2012).

83. Watanabe, T. K. et al. cDNA cloning and characterization of a human proteasomal modulator subunit, p27 (PSMD9). Genomics 50, 241–250 (1998).

84. Hori, T. et al. cDNA cloning and functional analysis of p28 (Nas6p) and p40.5 (Nas7p), two novel regulatory subunits of the 26S proteasome. Gene vol. 216 113–122 Preprint at https://doi.org/10.1016/s0378-1119(98)00309-6 (1998).

85. Verma, R. et al. Proteasomal proteomics: identification of nucleotide-sensitive proteasome-interacting proteins by mass spectrometric analysis of affinity-purified proteasomes. Mol. Biol. Cell 11, 3425–3439 (2000).

86. Dawson, Apcher, Mee & Mayer. Gankyrin is an ankyrin-repeat oncoprotein that interacts with CDK4 kinase and the S6 ATPase of the 26 S proteasome. Boll. Soc. Ital. Biol. Sper.

87. Park, Y. et al. Proteasomal ATPase-associated factor 1 negatively regulates proteasome activity by interacting with proteasomal ATPases. Mol. Cell. Biol. 25, 3842–3853 (2005).

88. Saeki, Y., Toh-E, A., Kudo, T., Kawamura, H. & Tanaka, K. Multiple proteasome-interacting proteins assist the assembly of the yeast 19S regulatory particle. Cell 137, 900–913 (2009).

89. Roelofs, J. et al. Chaperone-mediated pathway of proteasome regulatory particle assembly. Nature 459, 861–865 (2009).

90. Goldberg, A. L., Kim, H. T., Lee, D. & Collins, G. A. Mechanisms That Activate 26S Proteasomes and Enhance Protein Degradation. Biomolecules 11, (2021).

91. Kim, H. T. & Goldberg, A. L. UBL domain of Usp14 and other proteins stimulates proteasome activities and protein degradation in cells. Proc. Natl. Acad. Sci. U. S. A. 115, E11642–E11650 (2018).

92. Akahane, T., Sahara, K., Yashiroda, H., Tanaka, K. & Murata, S. Involvement of Bag6 and the TRC pathway in proteasome assembly. Nat. Commun. 4, 2234 (2013).

93. Guo, X. et al. UBLCP1 is a 26S proteasome phosphatase that regulates nuclear proteasome activity. Proc. Natl. Acad. Sci. U. S. A. 108, 18649–18654 (2011).

94. Hu, C., Tao, L., Cao, X. & Chen, L. The solute carrier transporters and the brain: Physiological and pharmacological implications. Asian J. Pharm. Sci. 15, 131–144 (2020).

95. Qian, F., Jiang, X., Chai, R. & Liu, D. The Roles of Solute Carriers in Auditory Function. Front. Genet. 13, 823049 (2022).

96. Murali Mahadevan, H., Hashemiaghdam, A., Ashrafi, G. & Harbauer, A. B. Mitochondria in Neuronal Health: From Energy Metabolism to Parkinson’s Disease. Adv Biol (Weinh) 5, e2100663 (2021).

97. Shrimanker, I. & Bhattarai, S. Electrolytes. in StatPearls (StatPearls Publishing, 2022).

98. Sheng, L., Leshchyns’ka, I. & Sytnyk, V. Neural cell adhesion molecule 2 promotes the formation of filopodia and neurite branching by inducing submembrane increases in Ca2+ levels. J. Neurosci. 35, 1739–1752 (2015).

99. Singh, V., Erady, C. & Balasubramanian, N. Cell-matrix adhesion controls Golgi organization and function through Arf1 activation in anchorage-dependent cells. J. Cell Sci. 131, (2018).

100. Haase, G. & Rabouille, C. Golgi Fragmentation in ALS Motor Neurons. New Mechanisms Targeting Microtubules, Tethers, and Transport Vesicles. Front. Neurosci. 9, 448 (2015).

101. Corradi, A. et al. SYN2 is an autism predisposing gene: loss-of-function mutations alter synaptic vesicle cycling and axon outgrowth. Hum. Mol. Genet. 23, 90–103 (2014).

102. Beggs, A. et al. Hypermethylation of SNAP91 as an alternative mechanism of epidermal growth factor signalling dysregulation: a genome-wide meta-analysis with validation of colorectal cancers. Lancet 383, S25 (2014).

103. Bleichert, F. & Baserga, S. J. Ribonucleoprotein multimers and their functions. Crit. Rev. Biochem. Mol. Biol. 45, 331–350 (2010).

104. Jiao, L. et al. Ribosome biogenesis in disease: new players and therapeutic targets. Signal Transduct Target Ther 8, 15 (2023).

105. Tye, B. W. et al. Proteotoxicity from aberrant ribosome biogenesis compromises cell fitness. Elife 8, (2019).

106. Butti, Z. & Patten, S. A. RNA Dysregulation in Amyotrophic Lateral Sclerosis. Front. Genet. 9, 712 (2018).

107. Gonçalves, I. do C. G., et al. Neuronal activity regulates DROSHA via autophagy in spinal muscular atrophy. Scientific Reports vol. 8 Preprint at https://doi.org/10.1038/s41598-018-26347-y (2018).

108. Barmada, S. J. et al. Autophagy induction enhances TDP43 turnover and survival in neuronal ALS models. Nat. Chem. Biol. 10, 677–685 (2014).

109. Giandomenico, S. L., Alvarez-Castelao, B. & Schuman, E. M. Proteostatic regulation in neuronal compartments. Trends Neurosci. 45, 41–52 (2022).

110. Dörrbaum, A. R., Kochen, L., Langer, J. D. & Schuman, E. M. Local and global influences on protein turnover in neurons and glia. Elife 7, (2018).

111. Kabashi, E., Agar, J. N., Strong, M. J. & Durham, H. D. Impaired proteasome function in sporadic amyotrophic lateral sclerosis. Amyotroph. Lateral Scler. 13, 367–371 (2012).

112. Ramachandran, K. V. & Margolis, S. S. A mammalian nervous-system-specific plasma membrane proteasome complex that modulates neuronal function. Nat. Struct. Mol. Biol. 24, 419–430 (2017).

113. Livnat-Levanon, N. et al. Reversible 26S proteasome disassembly upon mitochondrial stress. Cell Rep. 7, 1371–1380 (2014).

114. Aiken, C. T., Kaake, R. M., Wang, X. & Huang, L. Oxidative stress-mediated regulation of proteasome complexes. Mol. Cell. Proteomics 10, R110.006924 (2011).

115. Kumar Deshmukh, F., Yaffe, D., Olshina, M. A., Ben-Nissan, G. & Sharon, M. The Contribution of the 20S Proteasome to Proteostasis. Biomolecules 9, (2019).

116. Yoshida, Y. et al. E3 ubiquitin ligase that recognizes sugar chains. Nature 418, 438–442 (2002).

117. Kumanomidou, T. et al. The Structural Differences between a Glycoprotein Specific F-Box Protein Fbs1 and Its Homologous Protein FBG3. PLoS One 10, e0140366 (2015).

118. Schweiger, S. et al. Resveratrol induces dephosphorylation of Tau by interfering with the MID1-PP2A complex. Sci. Rep. 7, 13753 (2017).

119. Heinz, A., Schilling, J., van Roon-Mom, W. & Krauß, S. The MID1 Protein: A Promising Therapeutic Target in Huntington’s Disease. Front. Genet. 12, 761714 (2021).

120. Kickstein, E. et al. Biguanide metformin acts on tau phosphorylation via mTOR/protein phosphatase 2A (PP2A) signaling. Proc. Natl. Acad. Sci. U. S. A. 107, 21830–21835 (2010).

121. Schweiger, S. et al. The E3 ubiquitin ligase MID1 catalyzes ubiquitination and cleavage of Fu. J. Biol. Chem. 289, 31805–31817 (2014).

122. Strong, M. J., Donison, N. S. & Volkening, K. Alterations in Tau Metabolism in ALS and ALS-FTSD. Front. Neurol. 11, (2020).

123. Sun, S. et al. Phosphatase UBLCP1 controls proteasome assembly. Open Biol. 7, (2017).

124. Collins, G. A. & Goldberg, A. L. Proteins containing ubiquitin-like (Ubl) domains not only bind to 26S proteasomes but also induce their activation. Proc. Natl. Acad. Sci. U. S. A. 117, 4664–4674 (2020).

125. Sisodia, S. S. Nuclear inclusions in glutamine repeat disorders: are they pernicious, coincidental, or beneficial? Cell vol. 95 1–4 (1998).

126. Ross, C. A. Polyglutamine pathogenesis: emergence of unifying mechanisms for Huntington’s disease and related disorders. Neuron 35, 819–822 (2002).

127. Taylor, J. P., Hardy, J. & Fischbeck, K. H. Toxic proteins in neurodegenerative disease. Science 296, 1991–1995 (2002).

128. Enam, C., Geffen, Y., Ravid, T. & Gardner, R. G. Protein Quality Control Degradation in the Nucleus. Annu. Rev. Biochem. 87, 725–749 (2018).

129. de Almeida, M. et al. AKIRIN2 controls the nuclear import of proteasomes in vertebrates. Nature 599, 491–496 (2021).

130. Guzder, S. N., Sung, P., Prakash, L. & Prakash, S. Affinity of yeast nucleotide excision repair factor 2, consisting of the Rad4 and Rad23 proteins, for ultraviolet damaged DNA. J. Biol. Chem. 273, 31541–31546 (1998).

131. Shuck, S. C., Short, E. A. & Turchi, J. J. Eukaryotic nucleotide excision repair: from understanding mechanisms to influencing biology. Cell Res. 18, 64–72 (2008).

132. Doss-Pepe, E. W., Stenroos, E. S., Johnson, W. G. & Madura, K. Ataxin-3 Interactions with Rad23 and Valosin-Containing Protein and Its Associations with Ubiquitin Chains and the Proteasome Are Consistent with a Role in Ubiquitin-Mediated Proteolysis. Molecular and Cellular Biology vol. 23 6469–6483 Preprint at https://doi.org/10.1128/mcb.23.18.6469-6483.2003 (2003).

133. Kim, I., Mi, K. & Rao, H. Multiple interactions of rad23 suggest a mechanism for ubiquitylated substrate delivery important in proteolysis. Mol. Biol. Cell 15, 3357–3365 (2004).

134. Hiyama, H. et al. Interaction of hHR23 with S5a. The ubiquitin-like domain of hHR23 mediates interaction with S5a subunit of 26 S proteasome. J. Biol. Chem. 274, 28019–28025 (1999).

135. Sokolova, V., Li, F., Polovin, G. & Park, S. Proteasome Activation is Mediated via a Functional Switch of the Rpt6 C-terminal Tail Following Chaperone-dependent Assembly. Sci. Rep. 5, 14909 (2015).

136. VerPlank, J. J. S., Tyrkalska, S. D., Fleming, A., Rubinsztein, D. C. & Goldberg, A. L. cGMP via PKG activates 26S proteasomes and enhances degradation of proteins, including ones that cause neurodegenerative diseases. Proc. Natl. Acad. Sci. U. S. A. 117, 14220–14230 (2020).

137. Dupré, A.-I., Haccard, O. & Jessus, C. The greatwall kinase is dominant over PKA in controlling the antagonistic function of ARPP19 in Xenopus oocytes. Cell Cycle 16, 1440–1452 (2017).

138. Kors, S., Geijtenbeek, K., Reits, E. & Schipper-Krom, S. Regulation of Proteasome Activity by (Post-)transcriptional Mechanisms. Front Mol Biosci 6, 48 (2019).

139. Um, J. W. et al. ASK1 negatively regulates the 26 S proteasome. J. Biol. Chem. 285, 36434–36446 (2010).

140. Takenaka, S., Fujisawa, T. & Ichijo, H. Apoptosis signal-regulating kinase 1 (ASK1) as a therapeutic target for neurological diseases. Expert Opin. Ther. Targets 24, 1061–1064 (2020).

141. Fujisawa, T. et al. The ASK1-specific inhibitors K811 and K812 prolong survival in a mouse model of amyotrophic lateral sclerosis. Hum. Mol. Genet. 25, 245–253 (2015).

142. Moon, S. L., Morisaki, T., Stasevich, T. J. & Parker, R. Coupling of translation quality control and mRNA targeting to stress granules. J. Cell Biol. 219, (2020).

143. Li, S. et al. Quality-control mechanisms targeting translationally stalled and C-terminally extended poly(GR) associated with ALS/FTD. Proc. Natl. Acad. Sci. U. S. A. 117, (2020).

144. Langmead, B. & Salzberg, S. L. Fast gapped-read alignment with Bowtie 2. Nat. Methods 9, 357–359 (2012).

145. Trapnell, C. et al. Differential gene and transcript expression analysis of RNA-seq experiments with TopHat and Cufflinks. Nat. Protoc. 7, 562–578 (2012).

146. Leek, J. T. & Storey, J. D. Capturing heterogeneity in gene expression studies by surrogate variable analysis. PLoS Genet. 3, 1724–1735 (2007).

147. DeCaprio, J. & Kohl, T. O. Differential Detergent Lysis of Cellular Fractions for Immunoprecipitation. Cold Spring Harbor Protocols vol. 2020 db.prot098582 Preprint at https://doi.org/10.1101/pdb.prot098582 (2020).

148. Barrett, T. et al. NCBI GEO: archive for functional genomics data sets--update. Nucleic Acids Res. 41, D991–5 (2013).

149. Edgar, R., Domrachev, M. & Lash, A. E. Gene Expression Omnibus: NCBI gene expression and hybridization array data repository. Nucleic Acids Res. 30, 207–210 (2002).

150. Vizcaíno, J. A. et al. 2016 update of the PRIDE database and its related tools. Nucleic Acids Res. 44, D447–56 (2016).

